# Systems analysis of metabolic responses to a mixed meal test in an obese cohort reveals links between tissue metabolism and the gut microbiota

**DOI:** 10.1101/2022.04.26.489057

**Authors:** Peishun Li, Boyang Ji, Dimitra Lappa, Abraham S Meijnikman, Lisa M. Olsson, Ömrüm Aydin, Sjoerd C. Bruin, Arnold van de Laar, Valentina Tremaroli, Hao Luo, Jun Geng, Kimberly A. Krautkramer, Annika Lundqvist, Hilde Herrema, Albert K. Groen, Victor E.A. Gerdes, Thue W. Schwartz, Fredrik Bäckhed, Max Nieuwdorp, Louise E. Olofsson, Jens Nielsen

## Abstract

Individuals with prediabetes and type 2 diabetes mellitus (T2DM) have poor ability to adapt to diet-triggered perturbations. We investigated global metabolic responses to a mixed meal test (MMT) in morbidly obese individuals with different diabetic status by performing plasma metabolomic profiling. Abnormal metabolism of carbohydrates, (branched-chain) amino acids, fatty acids and acylcholines in individuals with (pre)diabetes was observed. Moreover, differences in metabolic responses were associated with altered fecal metagenomics and transcriptomes of liver, jejunum and adipose tissues, which revealed a modified gut microbiome and multi-tissue metabolism in individuals having insulin resistance. Finally, using integrative machine learning models, we built a predictive model based on metabolomics data after 2h MMT, and identified possible new biomarkers for glycemic control including N−acetylaspartate and phenylalanine-derived metabolites that may be useful for diagnosis, intervention and prevention of T2DM.

## Introduction

Type 2 diabetes mellitus (T2DM), characterized by hyperglycaemia, is one of the fastest increasing diseases worldwide^1–3^. Before individuals develop T2DM, they almost always have prediabetes (Pre-D). 5-10% of all individuals with Pre-D will annually progress to T2DM, and ∼70% will eventually develop T2DM over the course of their lifetime^4^. These individuals are characterized by higher than normal blood glucose levels that have not yet reached the threshold for diabetes diagnosis^4, 5^. Individuals with Pre-D and T2D not only manifest metabolic disorders at fasting, but also have a reduced ability to adapt to diet-triggered perturbations, e.g., the limited control for postprandial glycemic level^6, 7^. Insulin resistance and pancreatic beta-cell dysfunction play important roles in the metabolic imbalance^8, 9^. Postprandial blood glucose control is important for diabetes management^10^, and mixed meal test (MMT) have been often used to assess postprandial responses of glucose and insulin^11–14^. Previous studies have revealed postprandial effects on the metabolism of Pre-D and T2DM patients^15–17^. In addition, early studies reported that postprandial glucose responses were predictable based on personal and microbial compositional features using machine learning models^18, 19^. However, few studies have taken a holistic view on how different underlying factors, including metabolisms of the gut microbiota and human host, contribute to abnormally metabolic responses to a MMT in individuals with (pre)diabetes.

Multi-omics profiling and data integrations have been widely applied in Pre-D and T2DM studies^20–23^. Especially, untargeted metabolomics technologies have provided an opportunity to investigate the global metabolic changes in populations with (pre)diabetes^20, 21, 24^, but the serum metabolome in previous studies has almost been examined at fasting condition. Several studies have shown that disorders of branched-chain amino acids (BCAAs) metabolism contribute to insulin resistance^25, 26^. Here, we applied a combination of metabolomics profiling and MMT to dynamically quantify metabolic processes in response to a MMT, which provides novel insights into metabolic imbalance in obese individuals with Pre-D and T2DM. In addition, transcriptomics studies have previously suggested potential mechanisms involved in the pathogenesis of Pre-D and T2DM^27–30^, but the existing studies have focused on single-tissue transcriptomic profiling, including subcutaneous adipose tissue, visceral adipose tissue and pancreatic islets. Here, we systematically analyzed gene expression profiles of different human tissues (liver, jejunum, mesenteric and subcutaneous adipose tissues), which enables us to identify differences in multi-tissue metabolisms of individuals with different diabetic status. Also, recent studies have shown that the gut microbiota has already significant alterations of metabolic capacity in Pre-D and correlates to T2DM progression^31–35^. Microbial metabolites, including BCAAs and histidine-derived imidazole propionate, have been demonstrated to be associated with insulin resistance^21, 36^. To gain further insights into associations between the gut microbiota and the metabolic responses of individuals, here shotgun metagenomics was used to determine the microbiota composition and potential microbiome functions.

Based on these obtained multi-omics data, we finally predicted glucose responses to a MMT. Interestingly, the predictive models trained with metabolomics data (especially after 2h MMT) showed best performance. By integrating different data sets, we were able to reveal how metabolic changes in different organs, including the gut microbiome, interplay and based on this identify a number of metabolites and gut microbial signatures that may serve as novel biomarkers of glycemic control.

## Results

### Study population

In the present study, we recruited 106 individuals with obesity (BMI≥30 kg/m^2^) scheduled for bariatric surgery and included in the BARIA cohort with either normal glucose tolerance (NGT, n=27), Pre-D (n = 57) or T2DM (n = 22) based on the American Diabetes Association criteria^5, 37, 38^ (**Fig. 1a**). Baseline characteristics are summarized in **Table 1**. Individuals with Pre-D and T2DM (47.3 ± 9.5 and 47.1 ± 10.1 years) were older (*P*<0.05) than those with NGT (41.3 ±10.0 years). Fasting glucose, HbA1c and triglycerides levels were, as expected, significantly higher (*P*<0.05) in the T2DM and Pre-D groups compared with the NGT group (**Table 1**). Similarly, individuals in the T2DM group had elevated insulin resistance index (HOMA2-IR) and decreased magnesium levels compared with the NGT and Pre-D groups (*P*<0.05). Additionally, individuals with T2DM used more medication, including insulin, metformin, thiazide and statin compared with NGT and Pre-D (*P*<0.05 by Fisheŕs Exact test; **Supplementary Fig. 1**). Furthermore, blood samples for the two-hour MMT and metabolomics profiling were drawn within three months before the bariatric surgery (**Fig. 1a** and **Supplementary Table 1**). Also, biopsies from liver, jejunum and adipose tissues, and fecal samples were collected on the day of the surgery.

**Figure 1:**
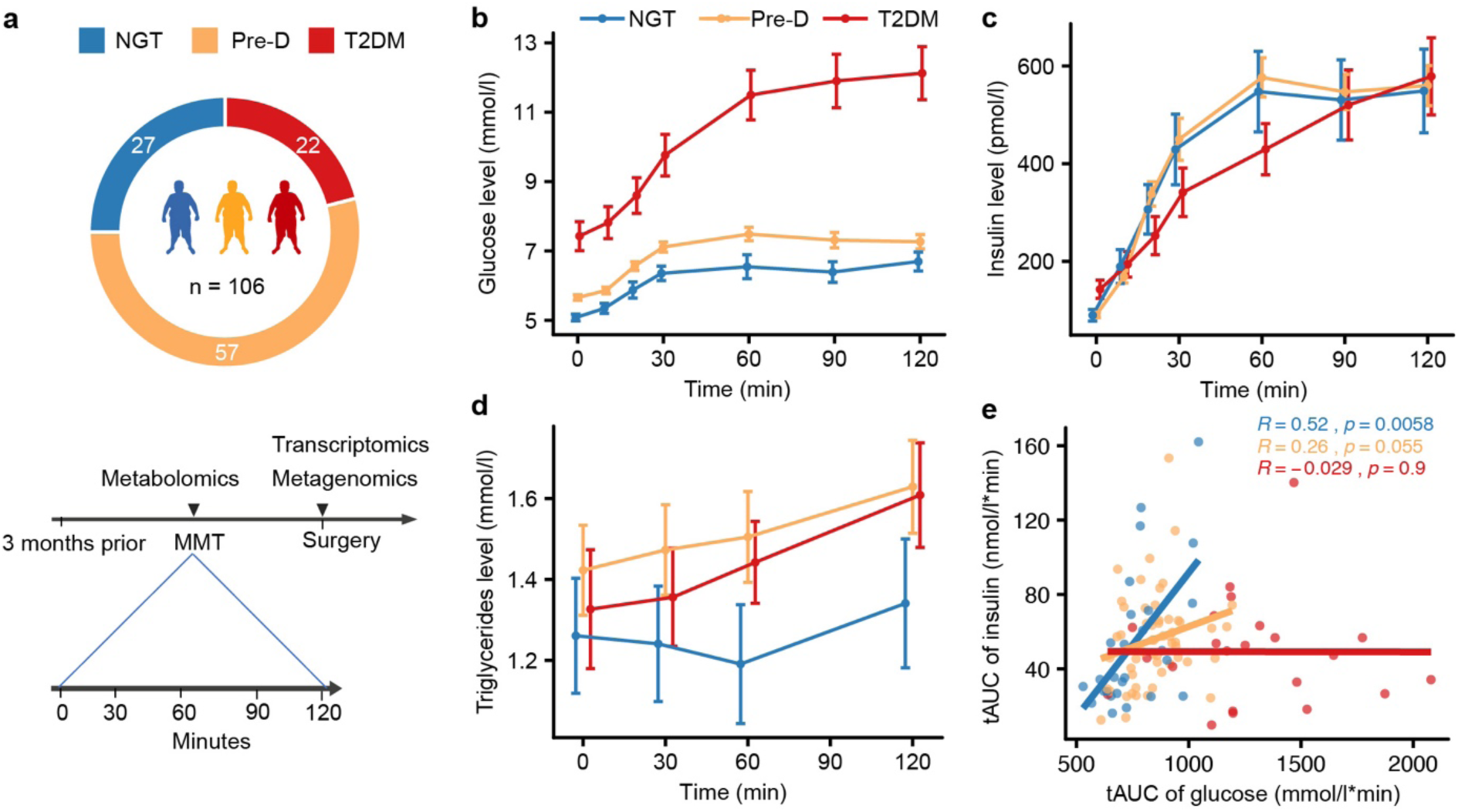
Experimental design and results from a mixed meal test (MMT). **a**, Schematic illustration of the experimental design. **b**, The time profiles of blood glucose, **c** insulin and **d** triglyceride concentrations during a MMT (Mean ± SEM) in the NGT (n=27), Pre-D (n=57) and T2DM (n=22). **e**, The association between insulin and glucose total AUC in each group. Spearman’s rank correlation analysis was performed.

**Table 1.** Baseline characteristics of the obese cohort. Mean±SD. For categorical variables number and percentages are presented. Non-normally distributed variables are presented as median with interquartile range. For comparison between groups, Fisheŕs Exact test was used for dichotomous variables and Student’s t-test or Wilcoxon rank sum test were used as appropriate for continuous variables. For comparison among three groups, Kruskal–Wallis test was used. ‘*’ denotes significant difference in comparison to NGT group (*P* < 0.05); ‘#’ denotes significant difference between Pre-D and T2DM groups (*P* < 0.05). BMI: body mass index, CRP: C-reactive protein, GGT: gamma glutamyl transferase, ALT: alanine aminotransferase, AST: aspartate aminotransferase, HbA1c: Hemoblobin A1c, HDL: high-density lipoprotein, LDL: low-density lipoprotein.

| Characteristics | NGT<br>(N = 27) | Pre-D<br>(N = 57) | T2DM<br>(N = 22) |
| --- | --- | --- | --- |
| **Demographic** |  |  |  |
| Age (years) | 41.3±10.0 | 47.3±9.5* | 47.1±10.1* |
| Female - no. (%) | 24 (88.9) | 44 (77.2) | 16 (72.7) |
| **Anthropometric** |  |  |  |
| BMI ($kg/m^2$) | 39.0 (37.5 to 40.9) | 39.8 (37.2 to 41.2) | 38.7 (35.9 to 42.7) |
| Weight (kg) | 113.0 (108.8 to 121.0) | 119.0 (104.8 to 128.0) | 118.5 (106.9 to 126.6) |
| Height (cm) | 170.0 (167.0 to 174.5) | 172.0 (167.0 to 178.0) | 172.5 (165.3 to 179.5) |
| Waist circumference (cm) | 118.0 (111.0 to 124.0) | 126.0 (117.8 to 135.0)* | 126.0 (117.0 to 130.0) |
| Total body fat (%) | 47.9 (46.8 to 50.1) | 47.6 (46.3 to 49.9) | 42.8 (40.1 to 48.5)* |
| Fat free mass (%) | 52.3 (50.0 to 52.3) | 52.5 (50.1 to 56.7) | 57.2 (51.5 to 59.9)* |
| Total body water (%) | 37.9 (35.6 to 39.9) | 39.4 (36.5 to 41.6) | 42.3 (38.5 to 46.0)*# |
| Systolic pressure (mmHg) | 132.0 (118.0 to 135.0) | 136.0 (126.0 to 143.0) | 130.0 (122.3 to 138.0) |
| Diastolic pressure (*mmHg*) | 80.0 (74.0 to 84.0) | 82.0 (79.0 to 88.0) | 83.0 (77.3 to 86.8) |
| **Clinical lab values** |  |  |  |
| Fasting glucose (*mmol/l*) | 5.1 (5.0 to 5.4) | 5.8 (5.6 to 6)* | 7.7 (6.8 to 8.9)*# |
| HbA1c (*mmol/mol*) | 34.0 (34.0 to 37.0) | 39.0 (37.0 to 40.0)* | 52.0 (46.0 to 60.0)*# |
| Fasting insulin (*pmol/l*) | 71 (45 to 116) | 80.5 (58.9 to 103.3) | 119.0 (64.3 to 212.3)*# |
| Fasting C-peptide (*pmol/l*) | 0.8±0.3 | 0.9±0.2 | 1.0±0.4 |
| Total cholesterol (*mmol/l*) | 4.6±1.0 | 5.2±1.1* | 4.3±1.0# |
| HDL-cholesterol (*mmol/l*) | 1.1 (1.0 to 1.4) | 1.2 (1.0 to 1.4) | 1.0 (0.9 to 1.2) |
| LDL-cholesterol (*mmol/l*) | 3.0±0.9 | 3.4±1.0 | 2.7±0.8# |
| Triglycerides (*mmol/l*) | 1.0 (0.8 to 1.4) | 1.4 (1.1 to 1.9)* | 1.6 (1.3 to 1.8)* |
| Ferritin ($\mu\text{g/l}$) | 70.0 (55.0 to 118.0) | 128.0 (56.0 to 184.0) | 85.5 (27.3 to 162.8) |
| CRP (*mg/l*) | 5.4 (3.4 to 10.5) | 4.9 (2.9 to 7.2) | 3.2 (5 to 9.4) |
| Hemoglobin (*mmol/l*) | 8.7 (8.1 to 9.0) | 9.0 (8.6 to 9.4) | 8.7 (8.4 to 9.2) |
| Leukocyte, 10<sup>9</sup>/l | 6.9 (6.1 to 8.8) | 6.9 (5.6 to 8.8) | 6.6 (5.6 to 9.4) |
| Creatinine ($\mu\text{mol/l}$) | 69.0 (61.0 to 78.0) | 66.0 (61.0 to 76.0) | 65.0 (56.8 to 76.3) |
| Magnesium (*mmol/l*) | 0.84±0.05 | 0.82±0.05 | 0.76±0.08*# |
| GGT (*IU/l*) | 22.0 (14.0 to 26.5) | 25.0 (18.0 to 35.3) | 38.0 (23.5 to 52.0)*# |
| ALT (*IU/l*) | 22.0 (19.5 to 30.5) | 27.0 (22.0 to 41.0) | 39.5 (27.0 to 56.0)*# |
| AST (*IU/l*) | 22.0 (20.0 to 25.5) | 24.0 (20.5 to 28.0) | 28.0 (21.3 to 40.8) |
| Vitamin B12 (*pmol/l*) | 321.0 (237.5 to 396.0) | 282.0 (219.0 to 360.0) | 263.0 (220.8 to 348.5) |
| Vitamin D (*nmol/l*) | 57.0 (31.5 to 71.5) | 48.0 (38.0 to 64.0) | 50.0 (30.0 to 71.0) |
| HOMA2-IR | 1.4 (0.8 to 2.2) | 1.6 (1.1 to 2.1) | 2.4 (1.5 to 4.5)*# |
| HOMA2-B | 129.8 (86.7 to 158.5) | 97.5 (84.0 to 120.7) | 81.0 (60.1 to 126.2)* |

### Mixed meal test characteristics

To assess the postprandial glucose response and insulin resistance, all individuals underwent a standardized MMT. The time profiles of blood glucose, insulin and triglyceride concentrations in responses to the MMT are shown in **Fig. 1b-d and Supplementary Fig. 2**. The MMT triggered a temporary increase in plasma glucose and insulin concentrations in NGT, Pre-D and T2DM groups (*P*<0.01 by ANOVA). However, glucose excursions differed significantly between the three groups (*P*<0.01 by ANOVA). This translated into significant differences in total area under the curve (tAUC) as well as in incremental AUC (iAUC, subtracting the baseline values) between the three groups (*P*<0.01 by Kruskal–Wallis test; **Supplementary Fig. 3**). T2DM individuals had slightly higher insulin levels at fasting, while had decreased insulin AUC/glucose AUC ratios and insulinogenesis index compared with NGT and Pre-D individuals (*P*<0.05 by Wilcoxon rank-sum test; **Supplementary Fig. 3**). The plasma triglyceride levels had an increasing trend over time in the Pre-D and T2DM groups, but with large within-group variations in the three groups (**Fig. 1d**). Overall, the responses of plasma glucose, insulin and triglyceride to the MMT were characterized by a prolonged elevation in the T2DM group. Interestingly, strong positive correlations between glucose AUC and insulin AUC were only observed in the NGT group (**Fig. 1e** and **Supplementary Fig. 4**; for tAUC, *R*=0.52 and *P*=0.0058; for iAUC, *R*=0.58 and *P*=0.0015). Thus, our results confirmed an abnormal plasma glucose and insulin response during a MMT for the Pre-D and T2DM groups.

### Metabolic signatures associated with diabetic status and modulated differentially by a MMT

To investigate the global metabolic responses to the MMT in individuals with different diabetic status, we used peripheral plasma samples for untargeted metabolomic profiling from participants at fasting and 2h post MMT and calculated the Euclidean distances between groups with different diabetic status at both time points (**Fig. 2a**). Interestingly, the three groups clustered separately at the two time points, which indicates that MMT has a large impact on the metabolomic profiles, with 439 metabolites being differentially abundant between the fasting and postprandial conditions (Adjusted *P*<0.05 by ANOVA; **Supplementary Table 2**). The main metabolic processes that had physiological responses to the MMT included lipids (n =183), amino acids (n=72) and xenobiotics (n=36) classes (**Supplementary Fig. 5**). Moreover, the distance between the NGT and T2DM groups at each time point shows an increased trend, compared to the distance between the NGT and Pre-D groups, or between the Pre-D and T2DM groups (**Fig. 2a**). This suggests a gradually increased difference in metabolomic profiles with progression from NGT to T2DM status. A total of 145 differential metabolites were associated with diabetic status (Adjusted *P*<0.05 by ANOVA, **Supplementary Table 2**), mainly composed of metabolites within the classes of lipids (n = 83), carbohydrates (n = 4), amino acids (n = 34), xenobiotics (n=6) and nucleotides (n = 7) (**Fig. 2b**). We further identified differentially abundant metabolites between any two groups by multiple pairwise comparisons at two time points, respectively (the bar chart in **Fig. 2b**; Adjusted *P* < 0.05 by t-test; **Supplementary Table 3**). No metabolites showed significantly different levels between the NGT and Pre-D groups at either time point. After 2h MMT, 139 metabolites differed between the NGT and T2DM groups, whereas only 55 metabolites differed at fasting. Consequently, metabolomic profiles were associated with diabetic status and showed a larger difference between the NGT and T2DM groups postprandially.

**Figure 2:**
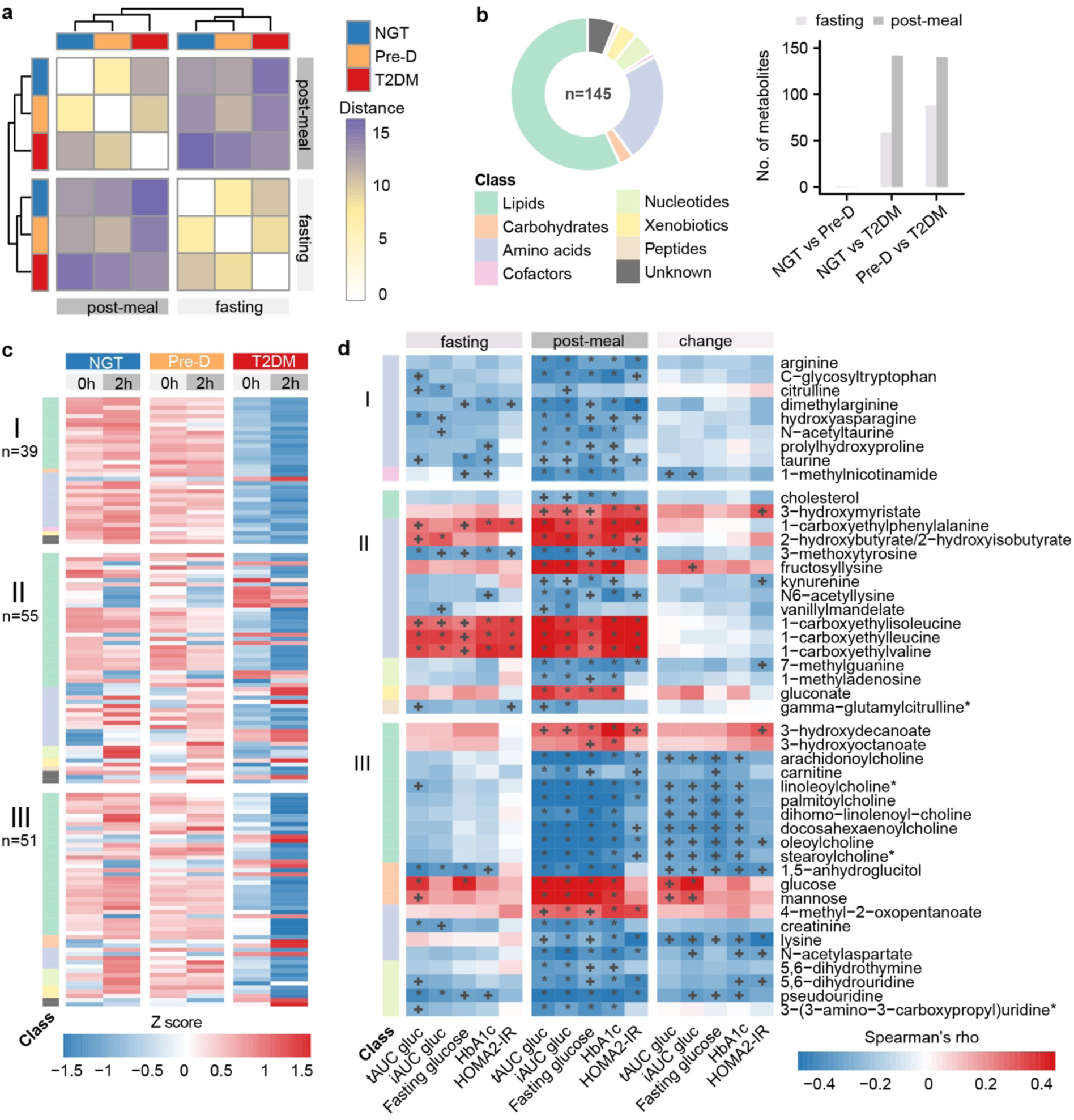
The metabolic changes associated with diabetic status and modulated differentially by the MMT. **a**, The hierarchical clustering of Euclidean distances between groups with different diabetic status at fasting and 2h post MMT. **b**, The donut chart shows pathways distribution of 145 metabolites differed significantly among the NGT (n=24), Pre-D (n=50) and T2DM (n=21) groups identified by ANOVA (adjusted *P* < 0.05). The bar chart shows the metabolites differed between the three groups at fasting and 2h post-meal, respectively. **c**, Heatmap showing the mean abundance of the metabolites with three different response patterns in the NGT (n=24), Pre-D (n=50) and T2DM (n=21) groups. **d**, The associations between the metabolomic changes and T2DM-related clinic variables. Only metabolites involved in the metabolic processes, including carbohydrates, amino acids, cofactors, nucleotides, xenobiotics, peptides, acylcholines, fatty acids, carnitine and sterol metabolism are shown. Spearman’s rank correlation analysis was performed. ‘+’ denotes adjusted *P* < 0.05; ‘*’ denotes adjusted *P* < 0.01. “biochemical name*” indicates a compound that has not been confirmed based on a standard but is confident in its identity.

Next, we classified the 145 metabolites associated with diabetes status into three response patterns, type I-III (**Fig. 2c**; **Method**; **Supplementary Fig. 6**; **Supplementary Table 3**). Of these, 39 metabolites showed a response pattern with no significant difference between the two time points in each group (type I) and 55 metabolites showed a parallel response to the MMT, independent of diabetes status (type II). The remaining 51 metabolites showed differential responses to the MMT among the three groups (type III) and primarily belonged to carbohydrates, amino acids and lipids. As expected, glucose abundance was significantly increased, whereas 1,5-anhydroglucitol abundance was decreased in the T2DM group compared to the NGT and Pre-D groups at both time points (adjusted *P* < 0.05; **Fig. 2c** and **Supplementary Table 3**). Moreover, mannose showed an elevated abundance in the T2DM group (adjusted *P* < 0.05), which is in agreement with previous results^39–41^. The identified metabolites in the carbohydrate class followed the type III response pattern, which indicates differential responses of carbohydrate metabolism to a MMT among the three groups.

Amino acid-derived metabolites including BCAAs and aromatic amino acids (AAAs) have been reported to be associated with T2DM^21, 36, 42^. Here, we identified 34 amino acid-derived metabolites associated with diabetes status (adjusted *P*<0.05 by ANOVA). Most amino acids in the type I response pattern, including arginine, taurine, N−acetyltaurine, C−glycosyltryptophan and hydroxyasparagine, showed reduced abundances in the T2DM group compared with the NGT and Pre-D groups at both time points (adjusted *P*<0.05; **Fig. 2c** and **Supplementary Table 3**). Additionally, amino acids in the type II response pattern, including 1−carboxyethylphenylalanine, 1−carboxyethylisoleucine, 1−carboxyethylleucine, 1−carboxyethylvaline and 2−hydroxybutyrate, were increased in the T2DM group compared to the NGT and Pre-D groups at both time points (adjusted *P*<0.05). Furthermore, amino acids with the type III pattern, including creatinine, lysine and N−acetylaspartate (NAA), were decreased in the T2DM group compared with the NGT and Pre-D groups at fasting or 2h post MMT (adjusted *P*<0.05). Particularly, these metabolites responded differentially to the MMT among the three groups, which may reflect the abnormal amino acid metabolism in the T2DM group after diet.

Many T2DM patients also have dyslipidemia^43^, and our analysis showed that abundances of metabolites in the fatty acid subclass were increased in the T2DM group at fasting or 2h post MMT (adjusted *P*<0.05, **Supplementary Table 3**). Six fatty acids, including 3−hydroxydecanoate, 3−hydroxyoctanoate, linoleate (18:2n6), linolenate (18:3n3 or 3n6), had differential responses to a MMT among the three groups (i.e., type III pattern). However, most metabolites in the lipid class had a decreased trend in the T2DM group compared to the NGT and Pre-D groups at both time points (**Fig. 2c** and **Supplementary Table 3)**, including sphingomyelin, carnitine, sterol, ceramides and acylcholine subclasses. Interestingly, all seven acylcholines, including arachidonoylcholine, linoleoylcholine*, palmitoylcholine, responded differentially to a MMT among the three groups.

### Associations of metabolomic changes with clinical variables

To assess the links between phenotypic characteristics and the postprandial changes of the T2DM-related circulating metabolites, we performed correlation analyses between the clinical variables and these metabolites at fasting or 2h post MMT (**Fig. 2d**). At fasting, most metabolites with type I and II response patterns had significant correlations with clinical variables glucose tAUC, HbA1c and HOMA2-IR (adjusted *P*<0.05, **Fig. 2d**). The carboxyethyl derivatives of BCAAs and phenylalanine were positively correlated with glucose tAUC and HOMA2-IR at both time points (adjusted *P*<0.05, *R*=0.29∼0.55). Interestingly, correlations between most metabolites and these clinical variables showed an increased trend after 2h MMT. For example, NAA only showed significant correlations with glucose AUC (both iAUC and tAUC) and HOMA2-IR after 2h MMT (adjusted *P*<0.05, for tAUC, *R*= -0.42; for iAUC, *R*= - 0.51; for HOMA2-IR, *R*= -0.36). Furthermore, correlations between the clinical variables and the MMT-induced metabolite changes (i.e., ratio of metabolite abundance at 2h post MMT to fasting) were investigated (**Fig. 2d**). The postprandial changes of most metabolites with type III pattern had significant correlations with the clinical variables (adjusted *P*<0.05). Especially, the postprandial changes of metabolites in acylcholine subclass were negatively correlated with glucose AUC (adjusted *P*<0.05, *R*= -0.4∼-0.33). In addition, the postprandial changes of NAA and lysine correlated negatively with HOMA2-IR (adjusted *P*<0.05, for NAA, *R*= -0.32; for lysine, *R*= -0.49), which suggests that the postprandial regulation of these metabolites might be associated with insulin resistance.

### Transcriptional changes associated with diabetes status

To identify differences in metabolic functions of individuals with variable diabetic status, gene expression profiles in four human tissues, including liver, jejunum, mesenteric and subcutaneous adipose tissue, were quantified using RNA-sequencing. As shown in **Fig. 3a,** the first and second principal component analysis (PCA) components clearly separated samples from different tissues, which accounted for 34% and 24% of the variability, respectively. But a few samples from mesenteric and subcutaneous adipose tissue overlapped. Differential gene expression analysis by multiple pairwise comparisons between the NGT, Pre-D and T2DM groups resulted in identification of 194, 30, 235 and 11 significantly differentially expressed genes in liver, jejunum, mesenteric and subcutaneous adipose tissue, respectively (adjusted *P*<0.05; **Supplementary Table 4**). These differentially expressed genes show tissue-specific **(Fig. 3b**). Furthermore, gene set analysis (GSA) identified enrichments of KEGG pathways in the four different tissues (*P*<0.05; **Fig. 3c**). Our results showed differences in metabolic pathway related to valine, leucine and isoleucine degradation in liver, jejunum and subcutaneous adipose tissue among the three groups. Differential genes involved in these pathways are summarized in **Supplementary Table 5** (*P*<0.01). In liver, insulin secretion, cAMP signaling pathway and cGMP−PKG signaling pathway were enriched with down-regulated genes, while steroid biosynthesis, terpenoid backbone biosynthesis and propanoate metabolism were enriched with up-regulated genes in the T2DM group compared with the NGT and Pre-D groups (*P*<0.05; **Fig. 3c**). The ryanodine receptor gene *RYR2* related to insulin secretion and genes related to cGMP-PKG signaling pathway including *IRS2*, *MYLK3*, *ADRA2C*, *NPPA,* were down-regulated in the T2DM group (*P*<0.01; **Supplementary Table 5**). In mesenteric adipose tissue, MAPK, TNF and NF-kappa B signaling pathway and cellular senescence were enriched with up-regulated genes in the T2DM group (*P*<0.05; **Fig. 3c**). In subcutaneous adipose tissue, the gene *NEU4* (encoding neuraminidase 4) related to sphingolipid metabolism was up-regulated in the Pre-D group compared with the NGT group (*P*<0.01 and |log_2_ (fold change)|>1). The gene *FASN* (encoding fatty acid synthase) involved in fatty acid metabolism was down-regulated in the T2DM group compared with the NGT group (*P*<0.01 and |log_2_ (fold change)|>0.6). In jejunum, genes *DGAT2, APOA4, MTTP, AGPAT2* related to fat digestion and absorption were up-regulated in the T2DM group compared with the Pre-D group (*P*<0.01; **Supplementary Table 5**).

**Figure 3:**
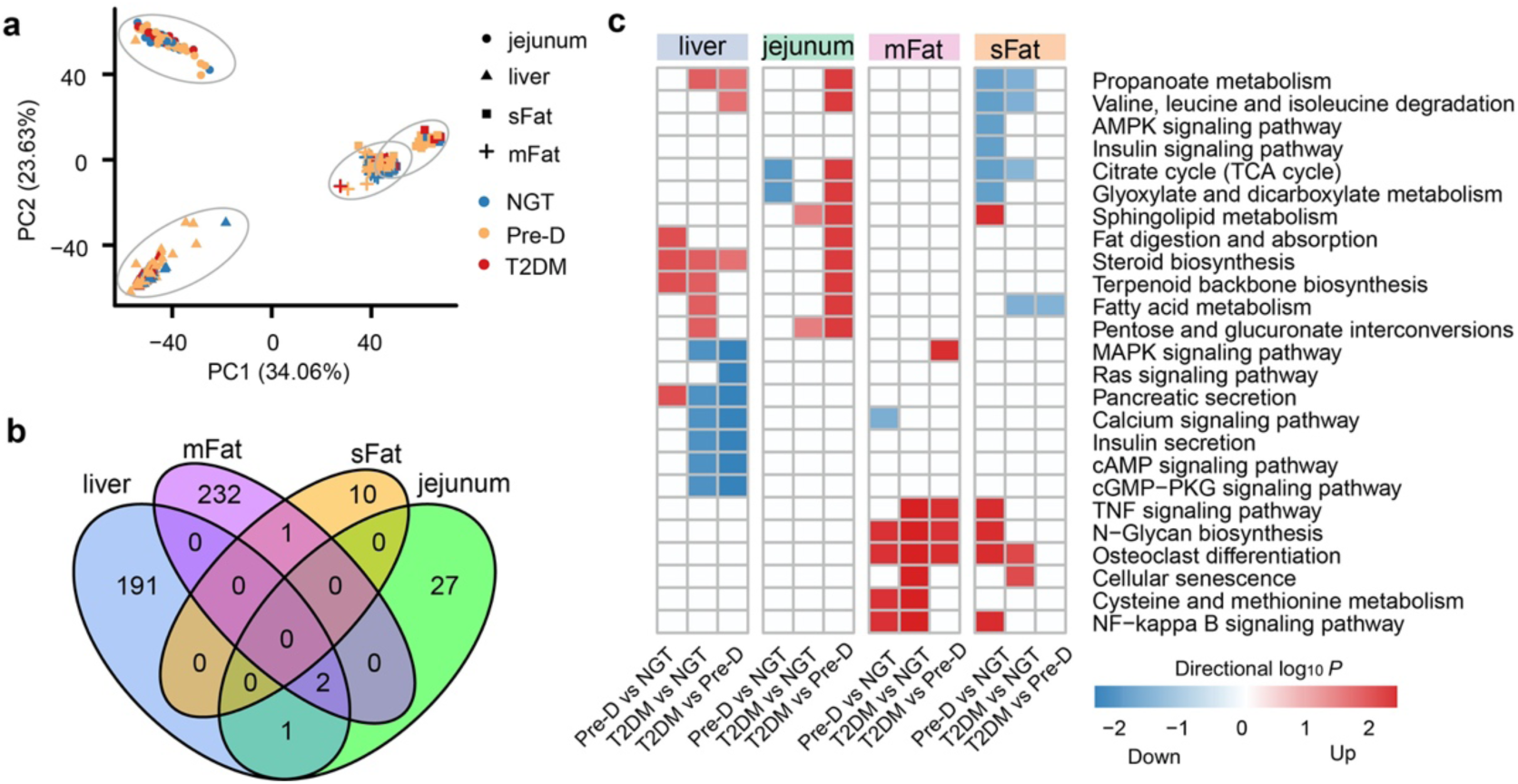
Transcriptional profiles of different human tissues from the NGT, Pre-D and T2DM individuals. **a**, Principle component analysis (PCA) of transcriptomic profiles in liver (n=106), jejunum (n=105), mesenteric(n=104) and subcutaneous adipose tissues (n=105). Nodes with circle, triangle, rectangle and crisscross represent samples from jejunum, subcutaneous and mesenteric adipose tissue, respectively. **b,** The Venn diagram depicting the distribution of significantly differentially expressed genes among the three groups in the four tissues (adjusted *P* < 0.05). **c,** The enriched KEGG pathways comparing the NGT, Pre-D and T2DM groups in the four tissues (*P* < 0.05). The red color indicates up-regulated gene sets; the blue color indicates down-regulated gene sets.

### Microbiota alterations associated with diabetes status

To determine the role of gut microbiota in the metabolic response to a MMT, the gut metagenome of the 106 individuals was quantified using shotgun DNA sequencing (**Supplementary Table 6**). Principal coordinate analysis (PCoA) shows that the second principal coordinate separates NGT and T2DM groups, which accounts for 11% of the variability (**Fig. 4a**). PERMANOVA analysis also shows that the diabetic status is associated with dissimilarities in gut microbiota composition (*R*^2^= 0.027, *P* <0.05). To identify differences in the bacterial composition among the three groups, microbial taxa at each taxonomical level, including class, order, family and genus, were compared by the Kruskal–Wallis test (**Supplementary Table 7**). Class Epsilonproteobacteria and from within this class, order Campylobacterales, family Campylobacteraceae and genus *Campylobacter* were more abundant in the Pre-D group than in the NGT and T2DM groups (adjusted *P* < 0.1 by Kruskal– Wallis test; **Fig. 4b** and **Supplementary Fig. 7**). In addition, family Peptostreptococcaceae with the genus *unclassified Peptostreptococcaceae* showed significantly decreased abundance in the T2DM group compared with the NGT and Pre-D groups (adjusted *P* < 0.1 by Kruskal–Wallis test; **Fig. 4b** and **Supplementary Fig. 7**), which is in accordance with a recent study^44^.

**Figure 4:**
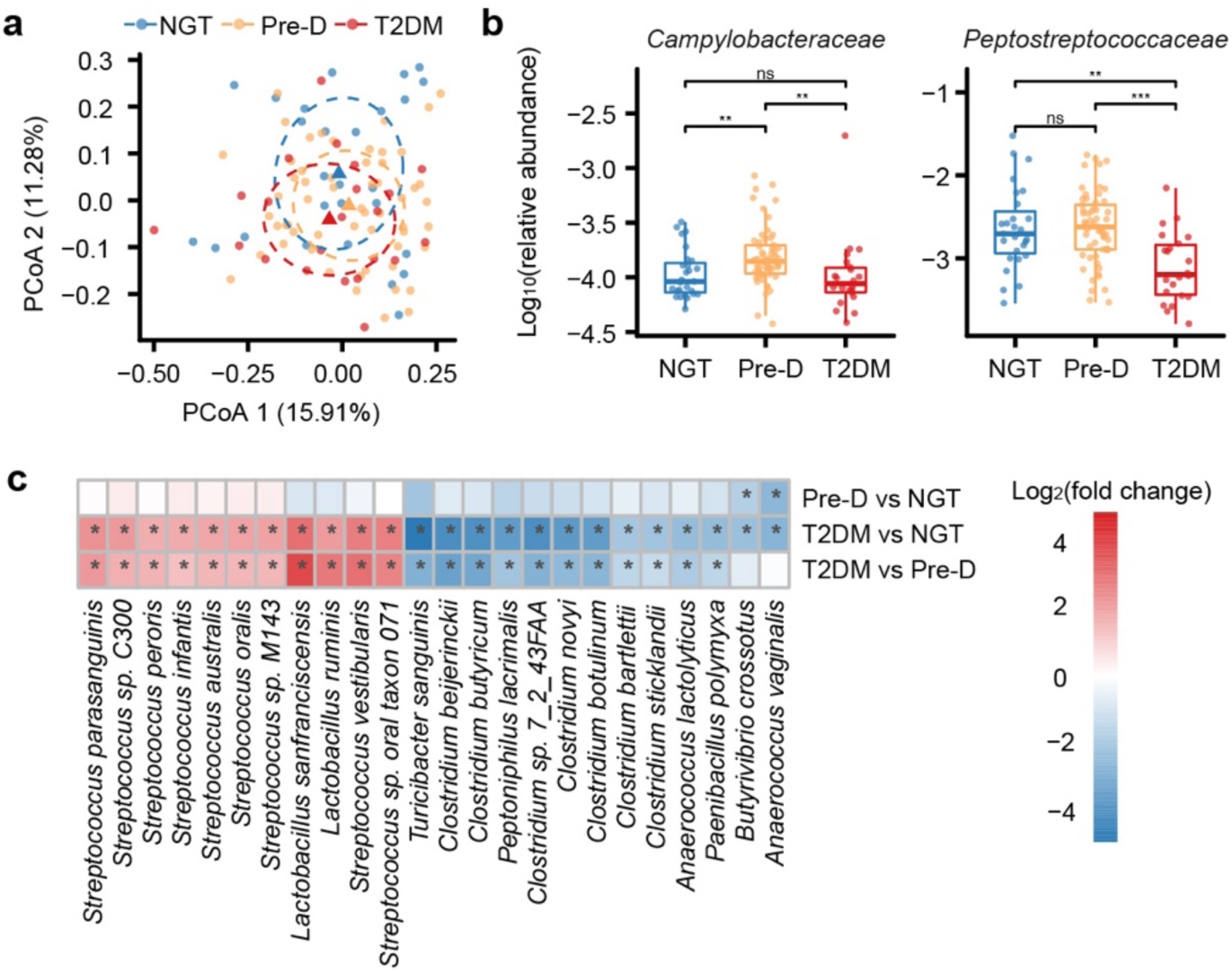
Alterations in gut microbiota associated with diabetes status. a, PCoA of microbiota community at species level based on Bray–Curtis distance (n=106). The centroid for each group is represented as a triangle and the ellipse covers the samples belonging to the group with 95% confidence. **b**, Log_10_ relative abundances of families Campylobacteraceae and Peptostreptococcaceae in the NGT (n=27), Pre-D (n=57) and T2DM (n=22), compared by Wilcoxon rank-sum test for multiple pairwise comparisons. ‘ns’ denotes no significance; ‘*’ denotes adjusted *P* < 0.05; ‘**’ denotes adjusted *P* < 0.01; ‘***’ denotes adjusted *P* < 0.001; ‘****’ denotes adjusted *P* < 0.0001. **c**, Heatmap showing log_2_ fold changes of 24 significantly differentially species between the NGT (n=27), Pre-D (n=57) and T2DM (n=22). Only species exhibiting differential abundance in two or three pairwise comparisons are shown. ‘+’ denotes adjusted *P* < 0.05; ‘*’ denotes adjusted *P* < 0.01.

Furthermore, 56 differential species were identified by multiple pairwise comparisons between NGT, Pre-D and T2DM groups (adjusted *P* <0.01; **Supplementary Table 8**), mainly belonging to the phylum Firmicutes. A total of 24 species exhibited differential abundance in two or three pairwise comparisons between the three groups (**Fig. 4c)**. The abundances of nine species of genus *Streptococcus*, *Lactobacillus sanfranciscensis* and *Lactobacillus ruminis* were increased, whereas the abundances of seven species of genus *Clostridium* (*Clostridium butyricum*, *Clostridium novyi*, etc.), *Turicibacter sanguinis*, *Anaerococcus lactolyticus* and *Paenibacillus polymyxa* were decreased in the T2DM group (adjusted *P* <0.01). Moreover, *Butyrivibrio crossotus* and *Anaerococcus vaginalis* were enriched in the Pre-D and T2DM groups compared to the NGT group (adjusted *P* <0.01). Especially, *C. novyi* had a significantly negative correlation with glucose iAUC (*R*=-0.49, *P* <1.0e-06; **Supplementary Fig. 8a**). Additionally, we investigated correlations between the differential species and the metabolites associated with diabetic status at fasting or 2h post MMT (**Supplementary Table 9**). *C. novyi* was positively correlated with NAA after 2h MMT (*R*= 0.46; *P* <1.0e-05; **Supplementary Fig. 8b**).

Interestingly, the correspondence between the gut microbiota composition and postprandial metabolomic profiles had an increased trend compared to the fasting condition (**Supplementary Fig. 9**). The carboxyethyl derivatives of BCAAs and phenylalanine were negatively correlated with several *Clostridium* species at both time points (*P*<0.01; **Supplementary Fig. 9**).

By investigating the functional capacity of the gut microbiome, we identified 60 significantly differential KOs among the NGT, Pre-D and T2DM groups (adjusted *P* <0.05; **Supplementary Table 10**). Moreover, we observed alterations in potential of phenylalanine and phenylacetate metabolism in the microbiome of individuals with Pre-D and T2DM by gene set analysis (*P*<0.05; **Table 2**). The microbial genes including *hcaC*, *hcaF*, *tynA*, *feaB, paaA* and *paaE* involved in phenylalanine metabolism, were more abundant in the T2DM group compared to the NGT or Pre-D group (*P*<0.01 and |log_2_ (fold change)|>3; **Table 2**). Especially, 1-carboxyethylphenylalanine correlated positively with genes *feaB, hcaC* and *paaE* (*P*<0.05 and *R*>0.2; **Supplementary Fig. 10**).

**Table 2.** The enriched KEGG pathways and modules in gut microbiome between the NGT (n=27), Pre-D (n=57) and T2DM (n=22) groups identified by gene set analysis. ’↑’ denotes significantly enriched pathway or module comparing two groups (*P* < 0.05). ‘–‘ denotes no differential genes in the pathway or module.

| | Differential genes ($P < 0.01$) | | |
| --- | --- | --- | --- |
|  | Pre-D vs NGT | T2DM vs NGT | T2DM vs Pre-D |
| **KEGG pathway** |  |  |  |
| Phenylalanine metabolism ↑ | – | *tynA, feaB, paaA, paaC, paaD, paaE, paaJ* | *hcaC, hcaF, paaJ* |
| **KEGG module** |  |  |  |
| Phenylacetate degradation ↑ | *paaE* | *paaA, paaC, paaD, paaE, paaJ* | – |

### Prediction of postprandial glucose response based on omics data

To systematically investigate potential contributing factors for metabolic responses to a MMT, we first quantified the associations between multi-omics data by the Mantel test using the Bray– Curtis distance (**Fig. 5a**; **Supplementary Table 11**). Significant correlations between the gut microbiome and metabolomics at fasting and 2h post MMT were observed (Mantel r = 0.10∼0.16, *P* < 0.05), which suggests that the gut microbiota is linked to metabolism of individuals in this study. Moreover, transcriptomics data correlated significantly in the different human tissues (Mantel r = 0.21∼0.28, *P* < 0.01). Interestingly, metabolomic changes (i.e., ratio of metabolite abundance at 2h post MMT to fasting) were correlated with the metabolomic profile at fasting (Mantel r = 0.084, *P* < 0.05), as well as transcriptional profiles in jejunum and mesenteric adipose tissue (Mantel r = 0.094 and 0.085, *P* < 0.05). This demonstrates that metabolic responses were associated with the metabolic profiles at baseline.

**Figure 5:**
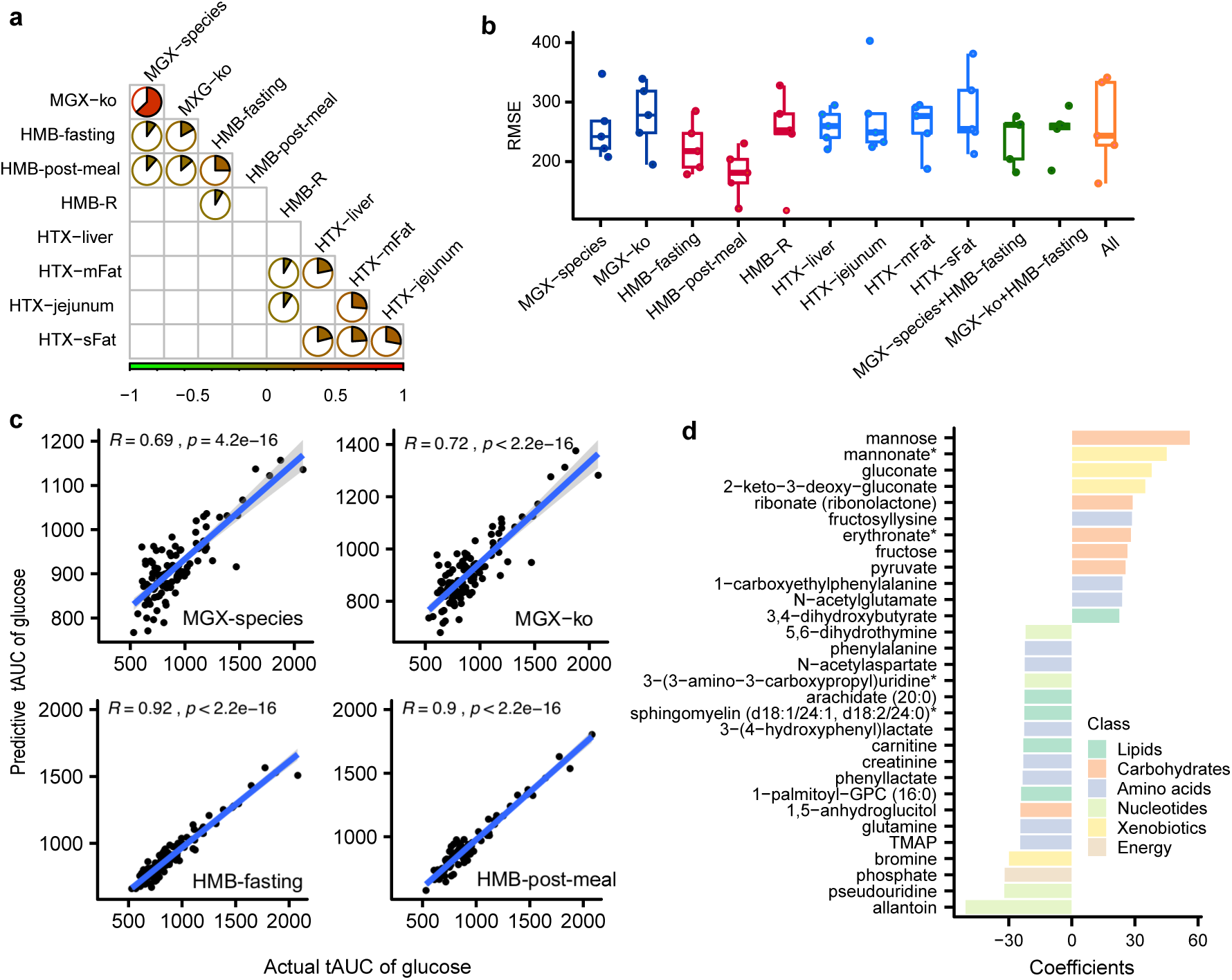
Predicting glucose response to a MMT by ridge regression models. **a**, The correlations between Bray–Curtis distance matrices of multi-omics. Mantel test was performed. The size of pie in the circle indicates the absolute value of correlation coefficient. Red and green colors represent positive and negative correlation coefficients, respectively. **b**, The performances of the ridge regression models evaluated by 5-fold cross-validation based on multi-omics and root mean square error (RMSE). **c**, The significant correlations between the actual glucose tAUC and the predicted glucose tAUC by ridge regression models using microbiota and metabolomics profiles, respectively. Spearman’s rank correlation analysis was performed. **d**, The regression coefficients of the top 30 metabolites for predicting glucose tAUC based on post-meal metabolomics data. MGX-species, microbiota composition at species level; MGX-ko, microbiota KO function profile; HMB-fasting, metabolomic profile at fasting; HMB-post-meal, metabolomic profile after 2h MMT; HMB-R, the ratios of metabolite abundance at 2h post MMT to fasting, which means the postprandial metabolic changes; HTX-liver, HTX-jejunum, HTX-mFat, HTX-sFat indicate human transcriptional profiles from liver, jejunum, mesenteric and subcutaneous adipose tissues, respectively; MGX-species+HMB-fasting, the combination of microbiota composition and metabolomic profile at fasting; MGX-ko+HMB-fasting, the combination of microbiota KOs profile and metabolomic profile at fasting; All, the integration of all multi-omics data.

To further investigate possible driving factors for postprandial glucose regulation, we predicted glucose tAUC based on multi-omics data using ridge regression models with 5-fold cross-validation. The models trained with metabolomics data (especially after 2h MMT) performed best with minimum root mean square error (RMSE) (**Fig. 5b**). The performance was improved when the model was trained using taxonomic (species) profiles compared to using functional (KOs) profiles and transcriptomics data. The correlation coefficients between the predictive and actual glucose tAUC were 0.92, 0.91 and 0.9 when using metabolomic profiles at fasting and 2h post MMT, and metabolomic changes as the training sets, respectively (**Fig. 5c** and **Supplementary Fig. 11**). Using species and KOs profiles of the gut microbiota as training sets, correlation coefficients between the predictive and actual glucose tAUC were 0.69 and 0.72, respectively (**Fig. 5c**). At fasting, glutamine, creatinine, pseudouridine, arginine, alanine, mannose, phenylalanine and lysine were identified to be the most important metabolites for prediction of glucose tAUC (**Supplementary Fig. 12a**). After 2h MMT, mannose, allantoin, phenylalanine, 1−carboxyethylphenylalanine and NAA were predicted to be the most important metabolites (**Fig. 5d**). Moreover, the postprandial changes of metabolites N-acetylalanine, carnitine, lysine, histidine, N-acetylserine and mannose were predicted to be important for glucose control (**Supplementary Fig. 12b**). The regression coefficients for species and KOs correlated with glucose tAUC are shown in **Supplementary Table 12**. We found that several *Clostridium* species, such as *Clostridium sp. D5, Clostridium sp. SS2* and *Clostridium bartlettii* were identified to be correlated with glucose tAUC. Consequently, our results revealed that glycemic response to a MMT was associated with the interaction of the gut microbiota and metabolism of individuals in this study.

## Discussion

Here, we revealed that the global metabolic responses to a MMT were different in individuals with varied glucose tolerance status. From plasma metabolomic profiling we found more differential metabolites between the NGT and T2DM groups after the meal intake compared to fasting condition, thus enabling us to discover abnormal metabolism related to (pre)diabetes that did not appear at fasting condition. Furthermore, we identified three different types of response patterns in the 145 metabolites that were associated with diabetic status. Following the MMT, 39 metabolites were unaltered; these were mainly amino acid-derived metabolites including arginine and taurine, that had reduced abundances in the T2DM group.

Another 55 metabolites showed a parallel response to the MMT in the NGT, Pre-D and T2DM groups, including the carboxyethyl derivatives of BCAA and phenylalanine that increased with elevating glucose level during the MMT. Also, these carboxyethyl derivatives were positively correlated with HOMA2-IR. Consistently, several studies in both rodents and humans have observed alterations in BCAA and amino acid metabolites in relation to insulin resistance^21, 26, 45^. Increase in circulating BCAA in cardiometabolic disease are considered to result from decreased catabolism in adipose tissue and from inactivation of the branched-chain ketoacid dehydrogenase (BCKDH) complex in the liver^26, 46, 47^. In line, we observed alterations in BCAA metabolism (i.e. valine, leucine and isoleucine degradation) in liver and subcutaneous adipose tissue by gene set analysis. Previous studies have suggested that the gut microbiome of individuals with insulin resistance also has an increased capacity to produce amino acids and specifically BCAA^21, 36^. By investigating the functional capacity of the gut microbiome, we also observed that amino acid metabolism (i.e. phenylalanine and phenylacetate metabolism) was enriched in the microbiome of individuals with insulin resistance. Interestingly, 1- carboxyethylphenylalanine correlated positively with microbial genes *feaB, hcaC* and *paaE* involved in phenylalanine metabolism. Microbial products of aromatic amino acid metabolism, in particular phenylacetic acid, has previously been linked to insulin resistance and thrombosis risk^48, 49^. Recently it was reported that phenylalanine-derived metabolites increased after autologous fecal microbiota transplantation (FMT) in individuals with liver steatosis^50^. Through integrative analysis, the carboxyethyl derivatives of BCAA and phenylalanine correlated negatively with several *Clostridium* species, indicating that a reduction of this bacterial species might influence changes in the circulating metabolites. Besides, Jagadeeshaprasad *et al* reported that quantification of 1−carboxyethyvaline peptides of beta-hemoglobin can be useful for assessing glycemic status^51^. Thus, these carboxyethyl derivatives of amino acids could be potential biomarkers for (pre)diabetes. Although our results are associative in nature, we further strengthen the hypothesis that the gut microbiome is capable of inducing alterations in circulating plasma metabolites. The strong correlations between circulating metabolites and insulin resistance, raise the question whether these metabolites serve as a biomarker or are causal agents in insulin resistance.

A total of 51 out of the 145 metabolites showed differential responses to the MMT in individuals with variable diabetic status. These metabolites might be involved in metabolic processes related to impaired adaptive response in (pre)diabetes, which commonly were not discovered at fasting. We observed different responses of carbohydrate metabolism (glucose, 1,5-anhydroglucitol, mannose) to the MMT, which is consistent with an earlier study^15^. Also, mannose has been identified as a biomarker of insulin resistance previously^40, 41^, which is in line with the fact that mannose is the most important metabolite for predicting glucose response in this study. Besides, several amino acids including N−acetylaspartate (NAA) responded differentially to the MMT. Interestingly, NAA involved in neuronal metabolism, was negatively correlated with HOMA2-IR, which is in agreement with a previous report^52^. NAA has been suggested to induce oxidative stress and nitric oxide (NO) production that has been reported to be associated with insulin resistance^53, 54^. Thus, NO synthesis may be down-regulated due to the decreased NAA abundance in individuals with insulin resistance. Meanwhile, the decreased arginine abundance may explain the reduced NO synthesis from arginine in the T2DM group^55^. In addition, reduced NO/cGMP signaling has been demonstrated to contribute to insulin resistance^56^. Consistently, the cGMP/PKG signaling pathway was down- regulated in the T2DM group in the liver, along with the decreased abundances of NAA and arginine. Through integrative analysis, *C. novyi* showed significantly positive correlation with NAA and negative correlation with glucose response. Thus, our results suggest potential interplay between *C. novyi*, NAA and insulin resistance via the NO/cGMP signaling pathway.

Additionally, fatty acids including 3−hydroxydecanoate, 3−hydroxyoctanoate, linoleate (18:2n6) and 3−hydroxysebacate, had a higher abundance in the T2DM group and showed differential responses to the MMT, which have previously been reported to be associated with insulin resistance^57–59^. Furthermore, the responses of acylcholines to the MMT were different in individuals with insulin resistance and negatively correlated with glucose response. Previous studies have reported that acylcholines can act as agonist of muscarinic acetylcholine receptors (mAChRs) and play an important role in stimulating insulin secretion and maintaining glucose homeostasis^60, 61^. Overall, the abnormal metabolism of carbohydrates, amino acids, fatty acids and acylcholines after a MMT in individuals with T2DM were revealed by metabolomic analysis in our study.

Interestingly, our results indicated that metabolic responses were associated with the metabolic status at baseline through integrative analysis. Therefore, differences in metabolic responses can be traced back to differences in other omics sets, such as liver, adipose tissue and jejunum transcriptomics data. By mapping genes onto KEGG pathways, we observed alterations in several pathways involved in crucial metabolic and inflammatory pathways in mesenteric adipose tissue, such as MAPK, TNF, NF-kappa B signaling and cellular senescence. The MAPK and NF-kappa signaling pathways have been suggested to be activated by TNF-alpha in adipose tissue, which is associated with insulin resistance^62–64^. Moreover, data from several human clinical studies has shown a clear correlation between insulin resistance and cellular senescence^65^. Furthermore, a previous study has suggested that RyR2 channels regulate insulin secretion and glucose homeostasis^66^. Our results also showed that the gene *RYR2* involved in insulin secretion was down-regulated in the T2DM group in liver, which is in accordance with the decreased HOMA2-B.

Several limitations of the current study must be acknowledged. Individuals with T2DM report a considerably higher number of medications than the NGT and Pre-D groups in this cohort, including glucose lowering agents, which may confound the T2DM-related changes in the gut microbiome and serum metabolomics (**Supplementary Table 13)**. In addition, the discrepancy in sampling time points may influence the results of our integrative analysis, but in this study it is no more than three months and has limited effect. To reduce the surgical risk, individuals have lost weight before the operation, which might introduce relevant biases in particular pre-operative weight-loss. However, in contrast to most bariatric surgery trajectories, these individuals did not adhere to a specific diet. Moreover, overfitting may happen due to the limited sample number and high number of features when we predicted glucose response using multi-omics data. Also, these identified contributing factors for postprandial glucose response need to be further validated in a new cohort. Another limitation is that issues associated with compositional data may impact the identified associations between metabolites and species^67^.

In conclusion, our study systematically characterized the metabolic response to a MMT in individuals with different glucose tolerance, which provides new insights into the metabolic imbalance of (pre)diabetes. We first identified the abnormal metabolic processes related to (pre)diabetes after meal intake, including carbohydrates, amino acids, fatty acids and acylcholines. Further, we revealed that differences in metabolic responses could be traced back to other omics sets including fecal metagenomics and transcriptomics data of liver, adipose tissue and jejunum. Using machine learning models, we identified possible new biomarkers for glycemic control including NAA and phenylalanine derived metabolites. However, future studies should test whether these potential biomarkers can be used for the early identification of individuals that are at risk of developing T2DM. Also, further studies are needed to validate the biological causality of the identified metabolic imbalance of (pre)diabetes.

## Methods

### Study population

We studied 106 individuals with morbid obesity in the BARIA cohort scheduled for bariatric surgery^37^. The study was performed in accordance with the Declaration of Helsinki and was approved by the Academic Medical Center Ethics Committee of the Amsterdam UMC. All participants provided written informed consent. Firstly, these individuals were classified into T2DM group (n=20) and non-T2DM group (n=86) according to the diagnosis information. The 86 non-T2DM individuals were further classified by their fasting blood glucose and HbA1c levels according to American Diabetes Association (ADA) criteria^5, 38^. 27 Individuals were classified into normal glucose tolerance (NGT) group having HbA1c level <39 mmol/mmol and fasting blood glucose level <5.6 mmol/L; 57 individuals were classified into prediabetes (Pre-D) group having HbA1c level 39–47 mmol/mmol or fasting blood glucose level 5.6–6.9 mmol/L; Two individuals were de novo classified into T2DM group having HbA1c level ≥48 mmol/mmol or fasting blood glucose level ≥7 mmol/L. Finally, the 106 individuals in the cohort were classified into three groups, including NGT (n=27), Pre-D (n=57) and T2DM group (n=22), for further analyses.

### Measurements of clinical characteristics

Individuals underwent a complete metabolic work-up at the start of their bariatric surgery trajectory. Anthropometric measurements including height, weight and waist circumference were taken. In addition, body fat percentage using bioelectrical impedance and blood pressure were measured. Fasting blood samples were used for the determination of fasting blood glucose, HbA1c, total cholesterol, HDL-cholesterol, LDL-cholesterol, triglycerides, ferritin, CRP (C-reactive protein), hemoglobin, leukocyte, creatinine, magnesium, GGT (gamma glutamyl transferase), ALT (alanine aminotransferase), AST (aspartate aminotransferase), vitamin B12, vitamin D and insulin levels.

### Mixed meal test

Within three months before surgery, individuals in the cohort underwent a 2-hour mixed meal test (MMT), which was performed to assess insulin resistance and investigate dynamic alterations in circulating metabolites. The MMT consisted of two Nutridrink compact 125ml (Nutricia®), containing 23.3 grams fat, 74.3 grams carbohydrates (of which 38.5 grams sugar) and 24.0 grams protein. The participants received this meal after fasting for a minimum of nine hours. Time point zero refers to the moment at which the participant had fully consumed the meal. Blood samples were drawn via an intravenous line at baseline, 10, 20, 30, 60, 90 and 120 minutes. Glucose, insulin and triglycerides were measured at these seven time points.

A number of variables related to insulin resistance, pancreatic β cell function and MMT were calculated using previously published methods^68^. The HOMA2 model (the updated HOMA model) was used to estimate insulin resistance (HOMA2-IR index) and pancreatic β cell function (HOMA2-B index) for an individual from fasting plasma glucose and fasting insulin concentrations measured in a MMT^69, 70^. To quantify the postprandial responses of glucose and insulin to a MMT, total area under the curve (tAUC) and incremental AUC (iAUC, subtracting the baseline values) were calculated from their measurements by the trapezoidal method^71^. For calculating the AUC, the k-Nearest Neighbors (KNN) method was performed for imputation of all missing values using R function knnImputation in DMwR with default parameters. Insulin AUC/glucose AUC ratios were calculated to estimate glucose-stimulated insulin secretion during 2h MMT. Besides, the insulinogenic index was calculated by dividing the insulin iAUC during the first 30 minutes by the glucose iAUC during the same period^72^.

### Metabolomics analysis

106 and 95 EDTA plasma samples were collected from participants at fasting and 2h after MMT, respectively. Samples were shipped to METABOLON (Morisville, NC, USA) for performing analysis using ultra high-performance liquid chromatography coupled to tandem mass spectrometry (LC-MS/MS) untargeted metabolomics, as previously described^36^. The metabolomic abundance obtained, underwent significant curation via metabolites’ pre-filtering, imputation for subsets of metabolites’ missing values and data normalization, in order to minimize the effect of artifacts in the downstream analysis. The abundances of all metabolites from fasting and post-meal samples were analyzed together in this study. The metabolomics dataset is comprised of 1345 metabolites with 1041 compounds of known identity (named metabolites) and 304 compounds of unknown structural identity (unnamed metabolites). Metabolomics prefiltering and imputation were performed by utilizing a variation of the Perseus platform^73^. Essentially, data has been pre-filtered so as to have a maximum of 25% missing values for a metabolite across all samples. This was followed by a log transformation of all the measured metabolites’ raw intensities across the entire dataset. Then, we calculated the total data mean and standard deviation (by omitting missing values). Taking into account that the metabolite intensities distribution is approximately following normality, we chose a small distribution 2.5 standard deviations away from the original data mean towards the left tail of the original data distribution, with 0.5 standard deviations width. This new shrunken range corresponds to the actual lowest level of detection by the spectrometer. Here by drawing random values from this mini distribution, we filled the missing prefiltered data of choice. Normalization was conducted to the total signal for each sample, since each sample is a separate injection on the mass spectrometer. Effective control for changes in sample matrix affects ionization efficiency, hence there can be inevitable differences in how much each sample is loaded onto the column with each injection, etc. Therefore, we summed up the total ion intensity (i.e. total signal) for each of the samples and identified the sample with the lowest total signal. After this we could proceed to calculating the correction factor for each sample:

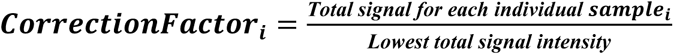

The next step is to divide each individual metabolite within a sample with the respective ***CorrectionFactor_i_***. Out of the 1345 metabolites analyzed by Metabolon, we used 998 metabolites in our downstream analysis after normalization and imputation.

To identify metabolites differential among the three groups with varied diabetic status or between two time points (fasting and 2h after MMT), two main effects (groups and time) and their interaction were assessed by multi-factor ANOVA that was adjusted for covariate age. Student’s t-test was used for multiple pairwise comparisons between two groups. *P* values were corrected for multiple testing using the false discovery rate method. Adjusted *P* < 0.05 was considered as threshold to identify significantly differential metabolites associated with diabetic status or the MMT. Due to the particular interest in (pre)diabetes-related metabolites’ responses to a MMT, the metabolites were further classified into three types of response patterns, as shown in **Supplementary Fig. 6**. Type I metabolites have no significant main effect for time and no interaction of two main effects time and groups, i.e., no response to a MMT. The first plot shows where the time profiles have no change and are parallel for the groups (parallel means no interaction). Type II metabolites have significant main effect for time but no interaction, i.e. parallel response to a MMT. The middle plots show where the time profiles have changes but are still parallel for the groups. Type III metabolites have significant interaction of two main effects, i.e., differential response to a MMT. The last plot shows where the time profiles have different changes for the three groups. The metabolites having differential responses among the three groups were identified by examining the significance of the interaction of two main effects (*P* <0.05). Moreover, to quantify the MMT-induced metabolic changes of each individual, the ratio of each metabolite abundance at 2h post MMT to fasting abundance was calculated.

### Transcriptome analysis

Biopsies from liver (106 samples), jejunum (105 samples), mesenteric adipose fat (104 samples) and subcutaneous adipose fat (105 samples) were collected at the time of the bariatric surgery (**Supplementary Table 1**), as previously described^37^. RNA was extracted from biopsies using TriPure Isolation Reagent (Roche) and Lysing Matrix D, 2 mL tubes (MP Biomedicals) in a FastPrep®-24 Instrument (MP Biomedicals) with homogenization for 20 seconds at 4.0 m/sec, with repeated bursts until no tissue was visible; homogenates were kept on ice for 5 minutes between homogenization bursts if multiple cycles were needed. RNA was purified with chloroform (Merck) in phase lock gel tubes (5PRIME) with centrifugations at 4°C, and further purified and concentrated using the RNeasy MinElute kit (QIAGEN, Hilden, Germany). The quality of RNA was analysed on a BioAnalyzer instrument (Agilent), with quantification on Nanodrop (Thermo Fisher Scientific). Due to degradation of the RNA, libraries for RNAseq sequencing were prepared by rRNA depletion; library preparation and sequencing were performed at Novogene (Nanjing, China) on an HiSeq instrument (Illumina Inc.) with 150 bp paired-end reads and 10G data/sample. The average read count per sample from liver and jejunum tissues are 42 ± 15 million. For mesenteric and subcutaneous adipose tissue, the average read count per sample are 43.2 ± 20 million.

Raw RNA-seq reads data were analyzed using nf-core/rnaseq^74^, a bioinformatics analysis pipeline for RNA sequencing data. Raw RNA-seq reads data was subjected to quality control using FastQC and multiQC^75^. The alignment of sequencing reads to the reference genome Homo sapiens GRCh38 was performed using STAR^76^. Gene counts were generated using featureCounts and StringTie^77, 78^. The pipeline was built using Nextflow^79^.

To identify the differential genes between NGT, Pre-D and T2DM groups, multivariate negative binomial generalized linear models were performed by R package DESeq2^80^. The models were adjusted for covariates age, BMI and gender. Only genes with the sum of counts across all samples ≥10 and existed in at least five samples were considered in the analysis. Raw read counts of genes were normalized using the median of ratios method by DESeq2. *P* values were corrected for multiple testing using BH method for per pairwise group comparison in each tissue. Adjusted *P*<0.05 was considered as threshold to identify significantly differentially expressed genes. To further explore differences in KEGG functions among the three groups, gene set analysis (GSA) was performed using statistics of all genes (*P* value and log_2_ fold change) and R package PIANO with the reporter algorithm for KEGG pathways^81^. The gene sets with a distinct directional *P* value<0.05 were chosen in this study, that is only considering gene sets significantly enriched by distinctly up or down-regulated genes.

### Microbiome analysis

Fecal samples from 106 participants were collected on the day of surgery and immediately frozen at -80C. Total fecal genomic DNA was extracted from 100 mg feces using a modification of the IHMS DNA extraction protocol Q^82^. Briefly, fecal samples were extracted in Lysing Matrix E tubes (MP Biomedicals) containing ASL buffer (QIAGEN), and lysis of cells was obtained, after homogenization by vortexing for 2 minutes, by two cycles of heating at 90°C for 10 minutes followed by three bursts of bead beating at 5.5 m/sec for 60 seconds in a FastPrep®-24 Instrument (MP Biomedicals). After each bead-beating burst, samples were placed on ice for 5 minutes. The supernatants containing fecal DNA were collected after the two cycles by centrifugation at 4°C. Supernatants from the two centrifugations steps were pooled and a 600 µL aliquot from each sample was purified using the QIAamp DNA Mini kit (QIAGEN) in the QIAcube (QIAGEN) instrument using the procedure for human DNA analysis. Samples were eluted in 200 µL of AE buffer (10 mmol/L Tris·Cl; 0.5 mmol/L EDTA; pH 9.0). Libraries for shotgun metagenomic sequencing were prepared by a PCR-free method; library preparation and sequencing were performed at Novogene (China) on an HiSeq instrument (Illumina Inc.) with 150 bp paired-end reads and 6G data/sample.

MEDUSA is an integrated pipeline for pre-processing of raw shotgun metagenomics sequence data^83^, which maps reads to reference databases, combines output from several sequencing runs and manipulates tables of read counts. The input number of total reads from the metagenome analysis were on average 23.4±2.2 million reads per sample and the total aligned reads were 16.6±1.8 million reads per sample (**Supplementary Table 6**). The sequencing runs had high quality with almost 98% of the reads passing the quality cut-off. Out of the high-quality reads, on average 0.04% aligned to the human genome, although the data had been cleaned for human reads. Out of the high quality non-human reads, 78.4% aligned to the MEDUSA’s software gene catalogue that contains more than 11 million genes^83^. Quality filtered reads were mapped to a genome catalogue that contains 1747 species genomes and the gene catalogue using Bowtie2^84^. Raw read counts at different taxonomy levels were normalized by scaling with cumulative sum (i.e. relative abundance). The α-diversity was calculated based on species-levels of each sample using Shannon, Simpson and Invsimpson indices via R package vegan^85^. To visualize and evaluate differences in gut microbiota composition among groups with varied diabetic status, principal coordinates analysis (PCoA) was performed based on species-level Bray–Curtis distances, and PERMANOVA was performed using the R function adonis in vegan. Microbial taxa at each taxonomical level, including class, order, family and genus, were compared by a Kruskal–Wallis test. *P* values were adjusted by FDR for each taxonomical level separately. To identify the differential species and KOs among the NGT, Pre-D and T2DM groups, multivariate negative binomial generalized linear models were performed by DESeq2 using raw read counts. The models were adjusted for covariates age, BMI and gender. *P* values were corrected for multiple testing using BH method for per pairwise group comparison. To further explore differences in KEGG functions among the three groups, gene set analysis (GSA) was performed using Piano with the reporter algorithm for KEGG pathways and modules. The differentially enriched KEGG pathways and modules were identified with a distinct directional *P* value<0.05.

### Statistical analysis

All statistical analyses were performed in the R software version 3.5. To identify the differential responses to a MMT, time curves of glucose, insulin and triglycerides concentrations for Pre-D and T2DM group were compared to NGT group using two-way ANOVA with repeated measures. The significant interaction of two main effects time (MMT) and groups (varied diabetic status) was investigated. In addition, Kruskal–Wallis test was used for comparisons among the three groups and Wilcoxon rank-sum test was used for multiple pairwise comparisons between each two groups. To assess the associations between multi-omics and clinical variables, Spearman’s rank correlation analysis was performed. *P* values were adjusted by FDR to control for multiple comparisons error in this study. To assess correlations between distance matrices of multi-omics, Mantel test was performed using the R package ade4 with the permutation number of 9999^86^. The Bray–Curtis dissimilarity matrices were calculated by using the R function vegdist in vegan.

To predict glucose response to a MMT (i.e. tAUC), ridge regression models were trained based on multi-omics data using R package glmnet^87^. The models were adjusted for covariates age, BMI and gender. First, ridge regression model was used to regress the normalized profile of gut microbiota, metabolomics, transcriptomics against glucose tAUC, respectively. Then, ridge regression model was used to regress the combinational normalized profile of multi-omics against glucose tAUC. The optimal lambda was chosen using function cv.glmnet (10-fold cross-validation in package glmnet) based on the minimum Root Mean Square Error (RMSE). To evaluate performance of the ridge model with the optimal lambda, 5-fold cross-validation (106 samples were randomly divided into five equal parts). Four fifths of samples were used to train the predicted model, and the remaining samples were used to test the fitness of it at each time) was performed by considering the measure RMSE.

## Acknowledgments

The BARIA study is funded by the Novo Nordisk Foundation (NNF15OC0016798). The Novo Nordisk Foundation Center for Basic Metabolic Research is supported by an unconditional grant (NNF10CC1016515) from the Novo Nordisk Foundation to University of Copenhagen. The BARIA study is a Scandinavian-Dutch collaboration. The computations and RNA Sequencing were enabled by resources provided by the Swedish National Infrastructure for Computing (SNIC) at C3SE (SNIC Computational Center of Chalmers University of Technology) partially funded by the Swedish Research Council through grant agreement no. 2018-05973. Funding from Knut and Alice Wallenberg Foundation is also acknowledged.

## Contributions

T.W.S., M.N., J.N. and F.B. supervised this work. P.L. and B.J. performed data analyses and visualization. D.L. processed metabolomics and transcriptomics data. L.M.O. processed metagenomics data. A.S.M., O.A., S.C.B. and A.V.D.L. collected medical data and biopsies. V.T. and A.L. performed DNA, RNA and metabolomics isolations and optimizations. H.L., J.G. and K.K. helped with ridge regression models, gut microbiota and metabolomics analysis. L.E.O., F.B. and J.N. coordinated project administration. P.L. wrote the first draft. P.L., B.J., A.S.M., L.E.O., H.H., A.K.G., V.E.A.G., T.W.S., M.N., F.B. and J.N. conducted hypothesis generation, manuscript review and editing.

## Availability of data and code

The related raw sequence files were submitted to public databases: the gut metagenomic data has been deposited in the European Nucleotide Archive (ENA) database under the accession number PRJEB47902; the transcriptomic data has been deposited in the European Genome-Phenome Archive (EGA) database with the access number EGAS00001005704, but the access to the transcriptomics data has been restricted to those who will get permission due to the GDPR. The metabolomic data has been deposited in the MetaboLights repository but can be accessed by contacting the leading authors with signing required documents. In addition, scripts used for the processing and analysis of data and other materials that support the findings of this study can be provided upon request.

## Supplementary Information

### Supplementary Figure Legends

**Supplementary Figure 1:**
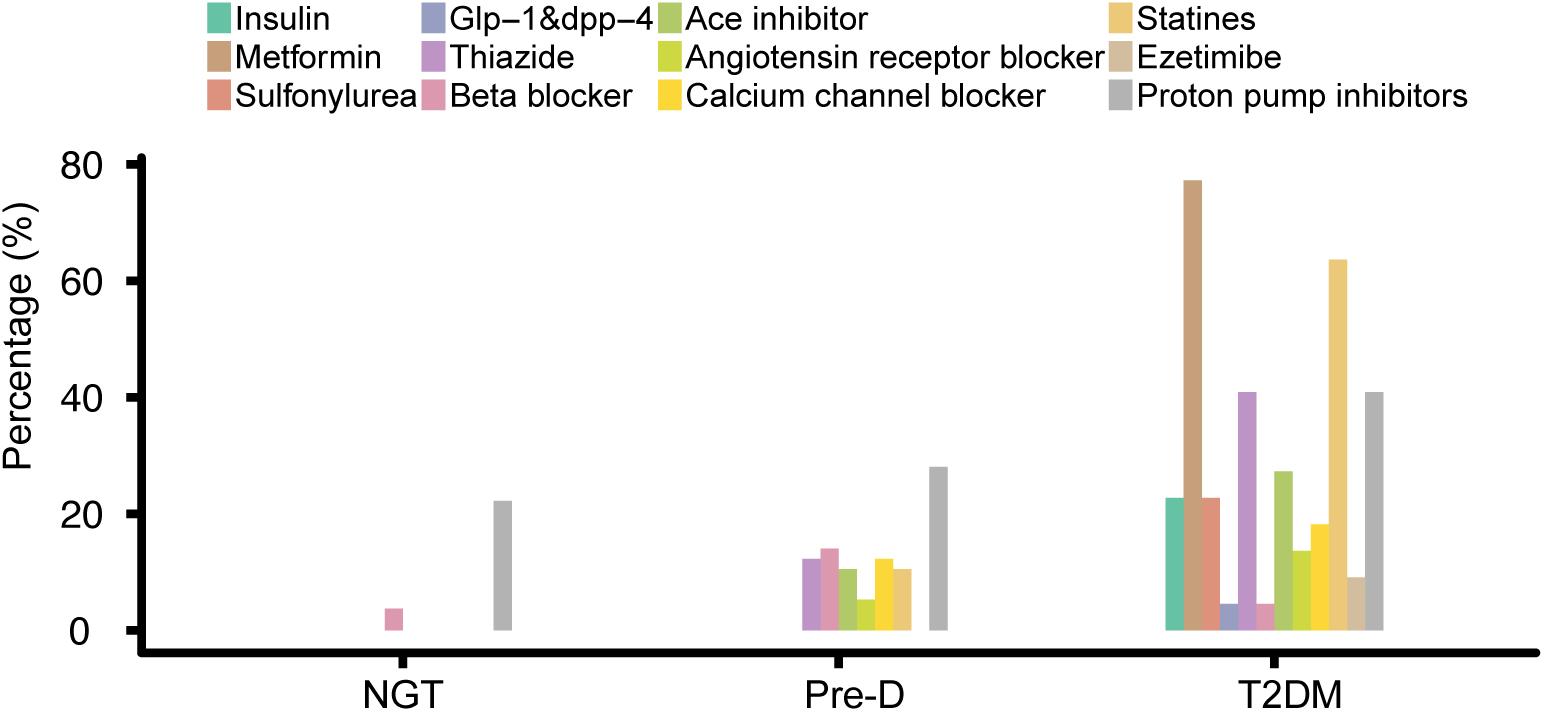
Comparisons of medication uses among NGT, Pre-D and T2DM groups. This bar chart shows that insulin, metformin, sulfonylurea, thiazide, ace inhibitor and statin use were correlated with diabetic status (*P* < 0.05). Fisheŕs Exact test was used for comparison among the three groups.

**Supplementary Figure 2:**
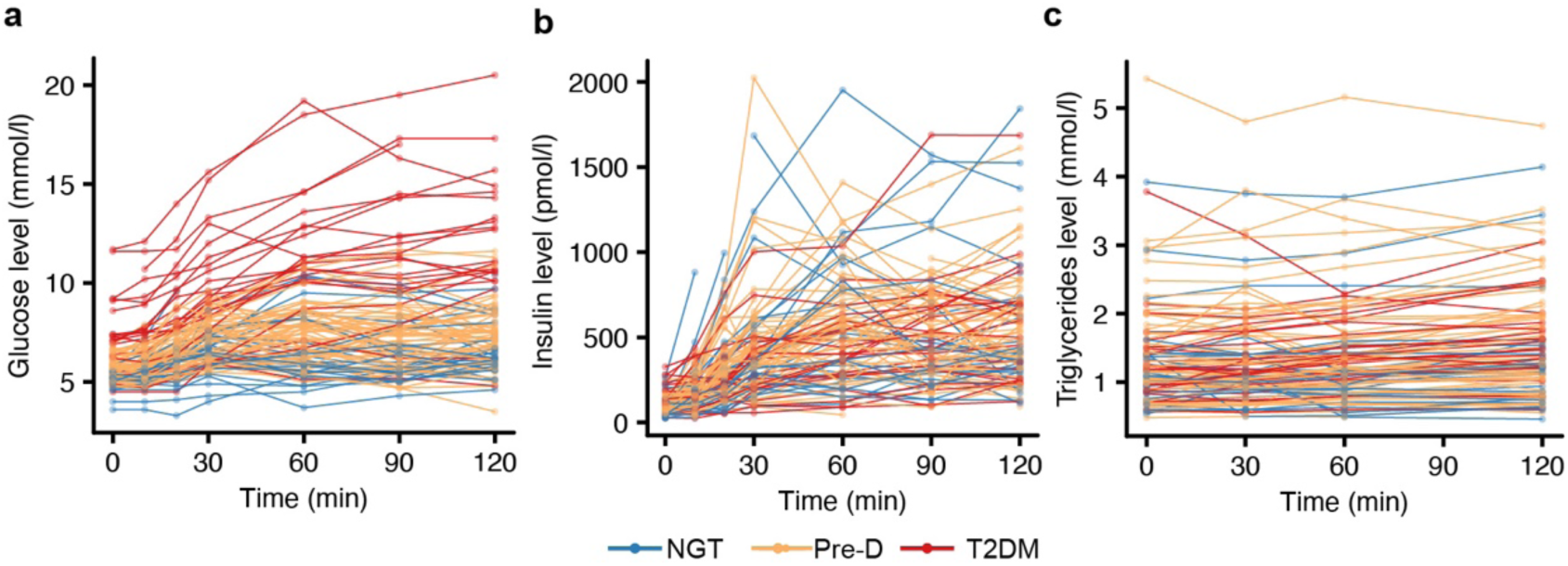
The individual time profiles of blood glucose (a), insulin (b) and triglyceride (d) concentrations during a 2h MMT.

**Supplementary Figure 3:**
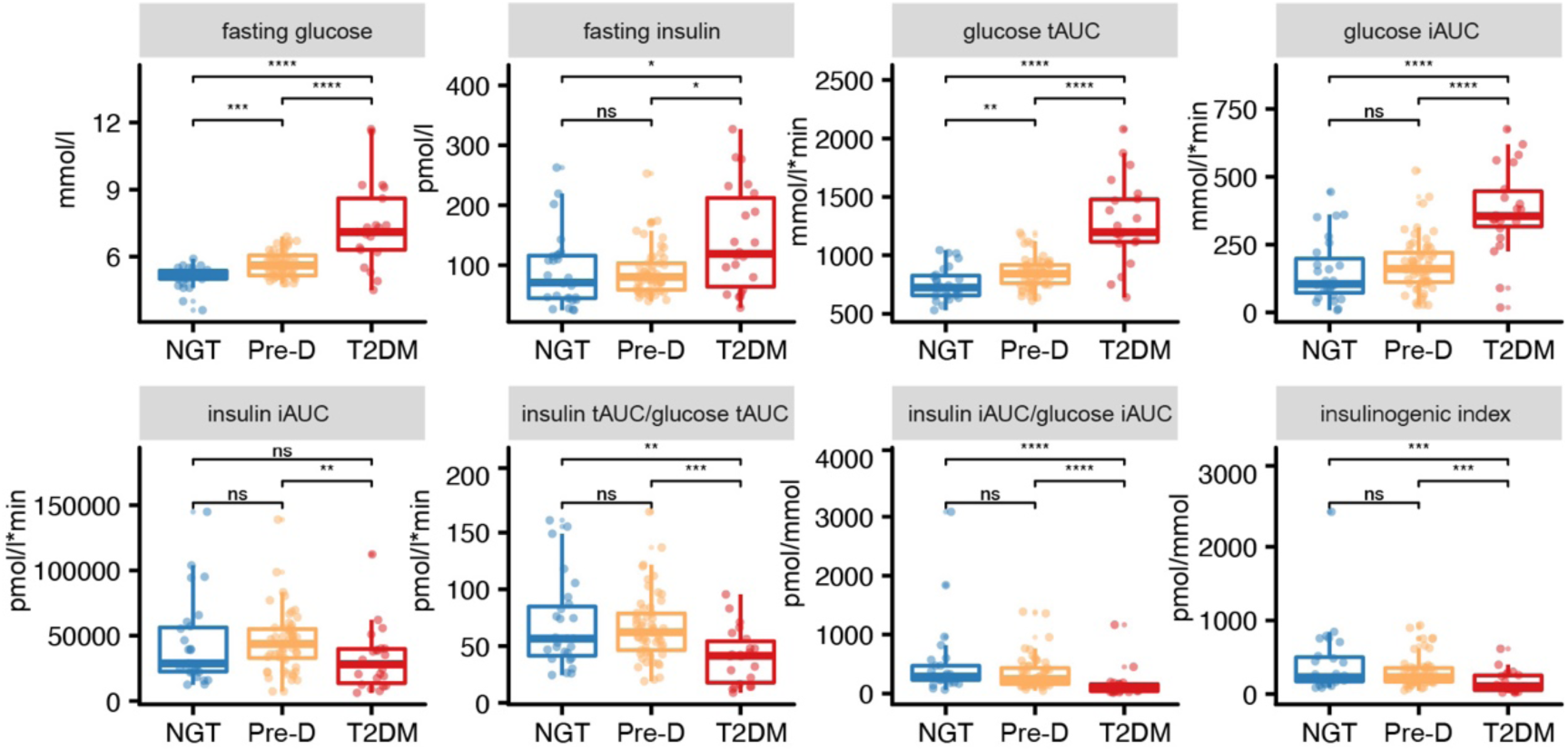
Comparisons of statistically significant characteristics related to the MMT. The differential characteristics were identified by Kruskal–Wallis test (*P* < 0.05) and compared by Wilcoxon rank-sum test for multiple pairwise comparisons. ‘ns’ denotes no significance; ‘*’ denotes *P* < 0.05; ‘**’ denotes *P* < 0.01; ‘***’ denotes *P* < 0.001; ‘****’ denotes *P* < 0.0001.

**Supplementary Figure 4:**
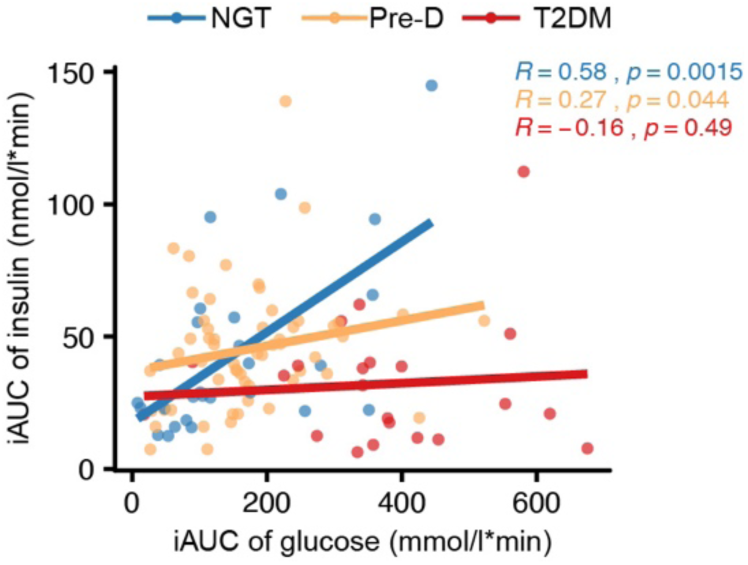
Significant association between insulin and glucose iAUC. Spearman’s correlation analysis was performed.

**Supplementary Figure 5:**
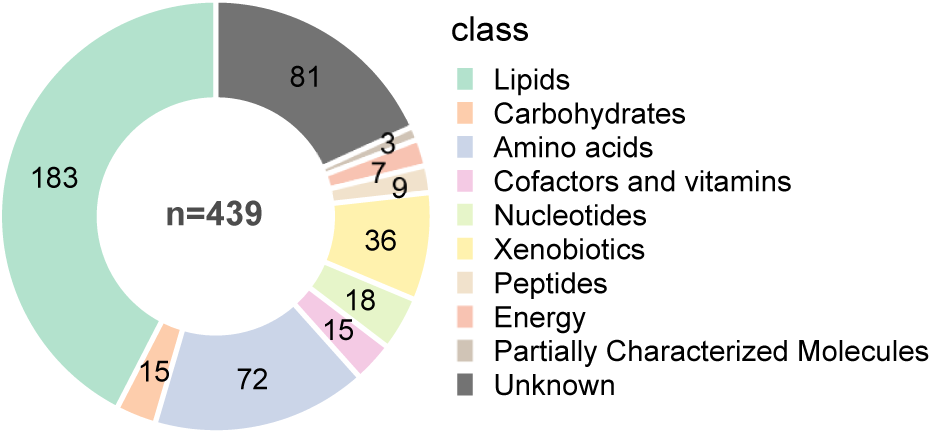
Metabolic processes that had physiological responses to a MMT. The significantly differential metabolites were identified by ANOVA (Adjusted *P* < 0.05).

**Supplementary Figure 6:**
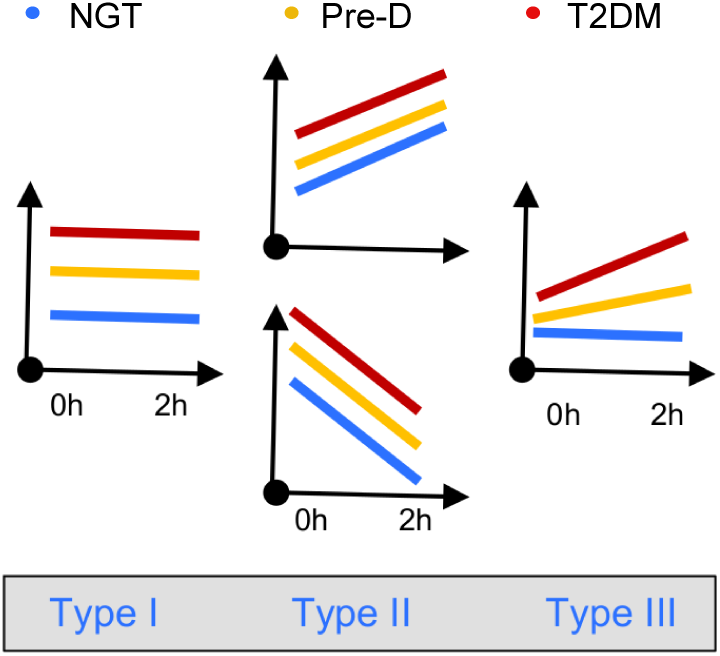
Schematic diagram showing three different types of metabolite response patterns. The first plot shows where the time profiles have no change and are parallel for the three groups (parallel means no interaction). The middle plots show where the time profiles have changes but are still parallel for the three groups. The last plot shows where the time profiles have different changes for the three groups.

**Supplementary Figure 7:**
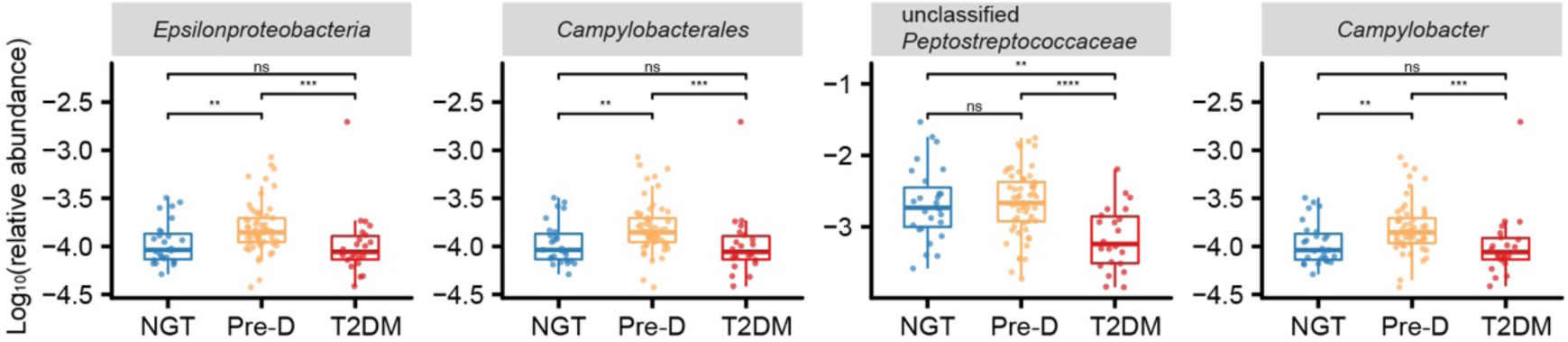
Taxonomical differences in bacterial composition. The abundances were compared by the Kruskal–Wallis test and by the Wilcoxon rank-sum test for multiple pairwise comparisons. ‘ns’ denotes no significance; ‘*’ denotes *P* < 0.05; ‘**’ denotes *P* < 0.01; ‘***’ denotes *P* < 0.001; ‘****’ denotes *P* < 0.0001.

**Supplementary Figure 8:**
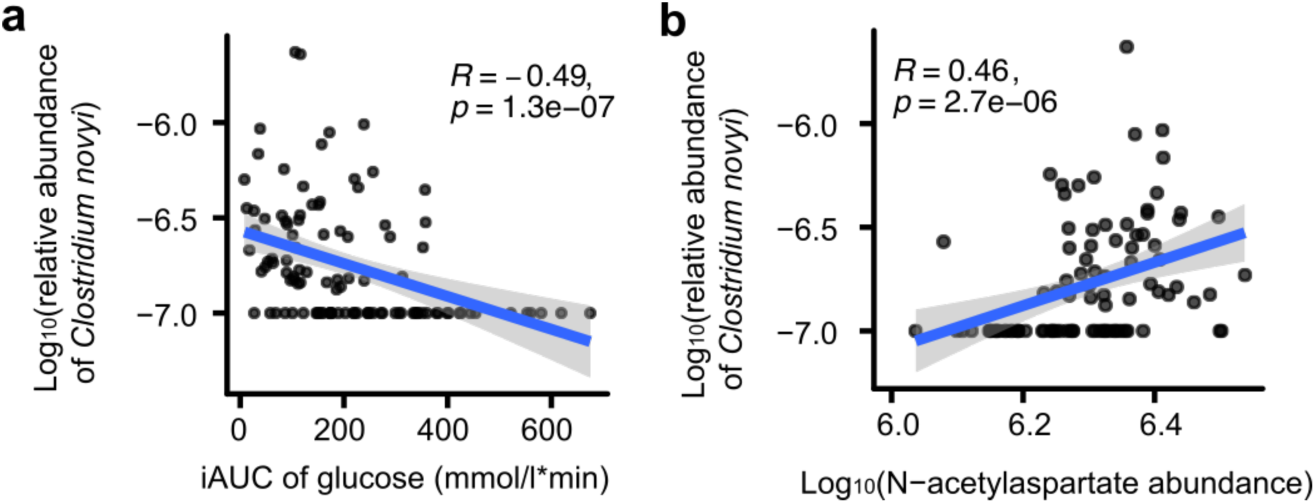
Significant associations of *Clostridium novyi* with (a) iAUC of glucose, and (b) N-acetylaspartate (NAA) after 2h MMT. Spearman’s rank correlation analysis was performed.

**Supplementary Figure 9:**
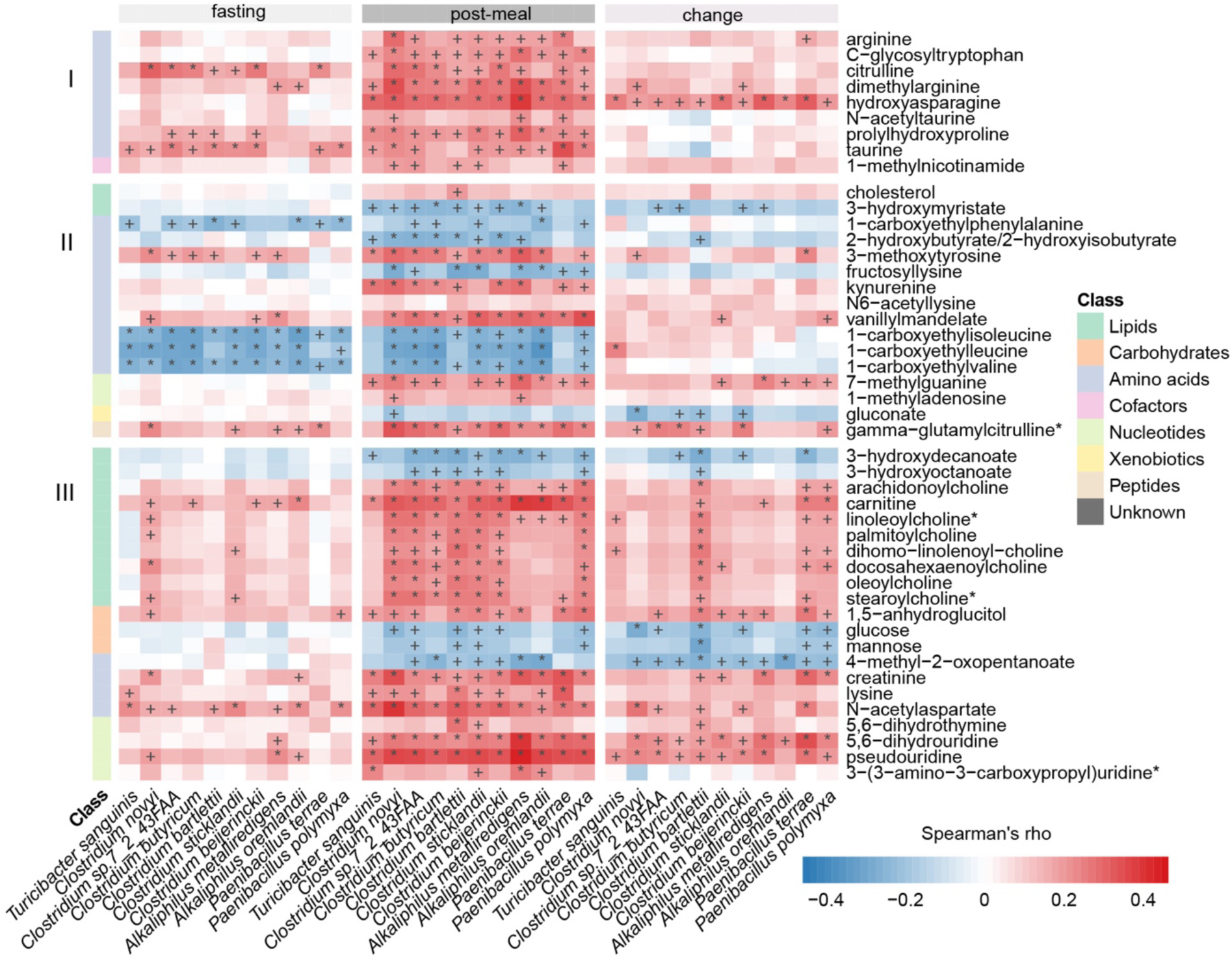
Associations between the metabolomic changes and the differential species. Only metabolites involved in the metabolic processes, such as carbohydrates, amino acids, cofactors, nucleotides, xenobiotics, peptides, acylcholines, fatty acids, carnitine and sterol metabolism are shown. Spearman’s rank correlation analysis was performed. ‘+’ denotes *P* < 0.05; The asterisk ‘*’ denotes *P* < 0.01.

**Supplementary Figure 10:**
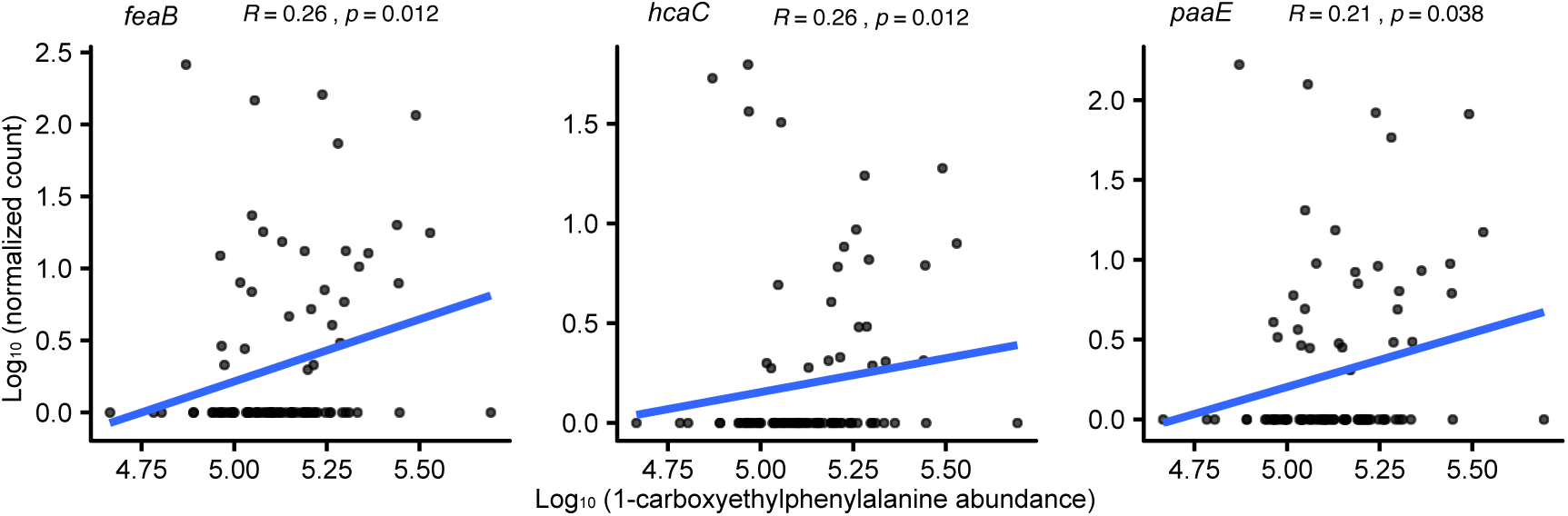
Associations between 1-carboxyethylphenylalanine and microbial genes *feaB*, *hcaC*, *paaE* involved in phenylalanine metabolism. Spearman’s rank correlation analysis was performed.

**Supplementary Figure 11:**
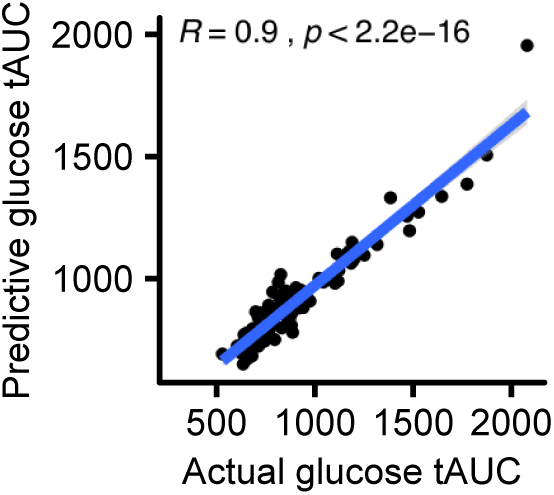
Strong correlation between the actual glucose tAUC and the predicted glucose tAUC by ridge regression model using metabolomic changes (i.e. ratio of metabolite abundance at 2h post MMT to fasting). Spearman’s rank correlation analysis was performed.

**Supplementary Figure 12:**
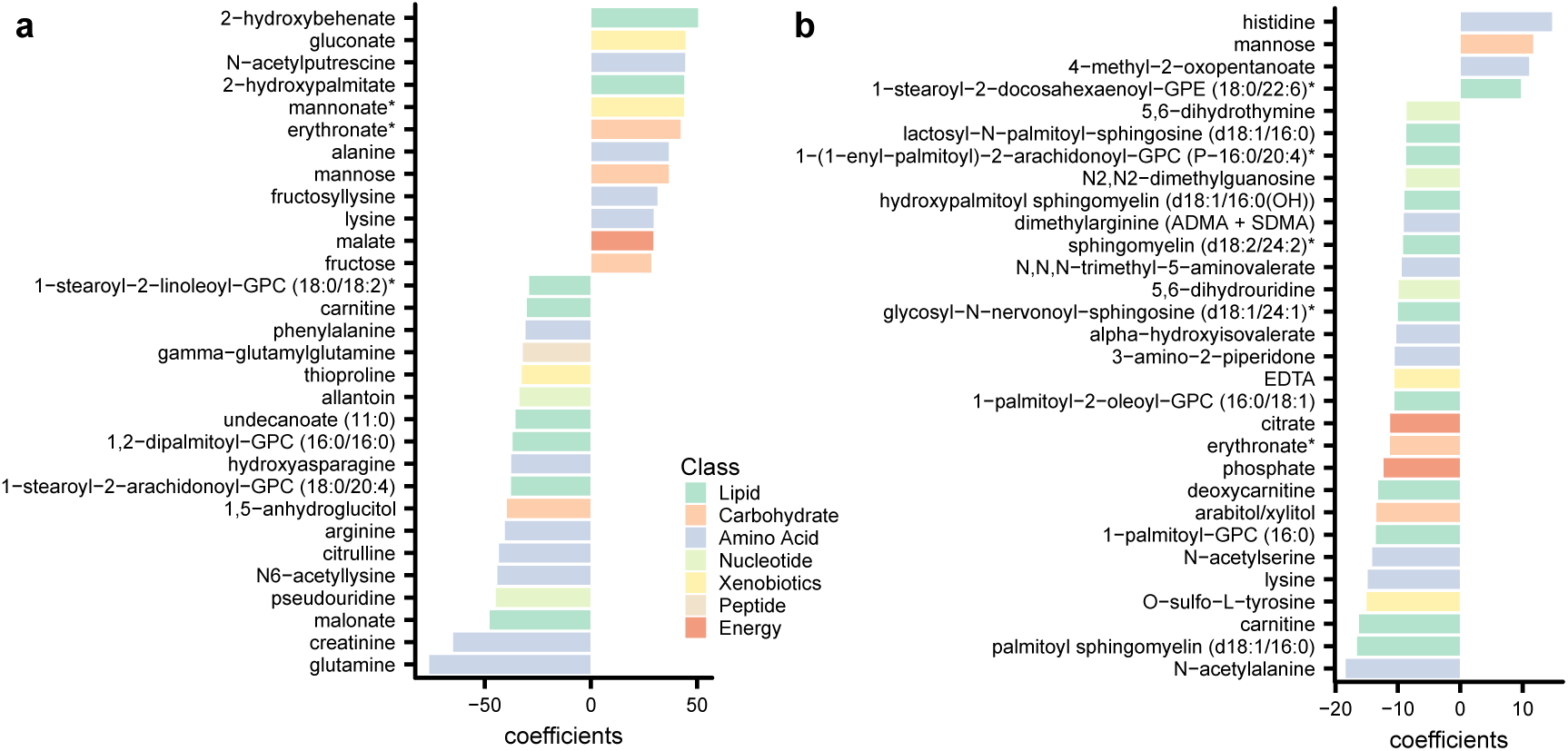
Regression coefficients of the top 30 important (a) metabolites at fasting, and (b) metabolites’ changes for prediction of glucose tAUC.

### Supplementary Table Legends

**Supplementary Table 1: Transcriptomic profiling of different human tissues and metagenomic profiling for 106 individuals in the BARIA cohort.**

**Supplementary Table 2a: Metabolites significantly affected by a MMT or associated with diabetic status identified by multi-factor ANOVA with adjustment for age.**

**Supplementary Table 2b: Metabolites significantly affected by a MMT or associated with diabetic status identified by two-way ANOVA.**

**Supplementary Table 3: Multiple pairwise comparisons of the 145 metabolites associated with diabetic status and the classifications of these metabolites into three different response patterns.**

**Supplementary Table 4a: Differential expressed genes associated with diabetes status in liver.**

**Supplementary Table 4b: Differential expressed genes associated with diabetes status in jejunum.**

**Supplementary Table 4c: Differential expressed genes associated with diabetes status in mesenteric adipose tissues.**

**Supplementary Table 4d: Differential expressed genes associated with diabetes status in subcutaneous adipose tissues.**

**Supplementary Table 5: The enriched KEGG pathways between NGT, Pre-D and T2DM groups in liver, jejunum, mesenteric and subcutaneous adipose tissues identified by gene set analysis and the differential genes involved in these pathways.**

**Supplementary Table 6: The mapping summary of metagenomic sequence files for 106 individuals in the BARIA cohort.**

**Supplementary Table 7a: Comparisons of microbial taxa at phylum level by Kruskal– Wallis test.**

**Supplementary Table 7b: Comparisons of microbial taxa at class level by Kruskal–Wallis test.**

**Supplementary Table 7c: Comparisons of microbial taxa at order level by Kruskal–Wallis test.**

**Supplementary Table 7d: Comparisons of microbial taxa at family level by Kruskal– Wallis test.**

**Supplementary Table 7e: Comparisons of microbial taxa at genus level by Kruskal–Wallis test.**

**Supplementary Table 8: Differential species among NGT, Pre-D and T2DM groups.**

**Supplementary Table 9: Correlation analysis between differential species and metabolites associated with diabetic status after 2h MMT.**

**Supplementary Table 10: Differential KOs among NGT, Pre-D and T2DM groups.**

**Supplementary Table 11: Associations between multi-omics data evaluated by Mantel test based on Bray–Curtis distance.**

**Supplementary Table 12a: Complete list of regression coefficients of models trained by metabolomic profiles at fasting.**

**Supplementary Table 12b: Complete list of regression coefficients of models trained by metabolomic profiles at 2h post MMT.**

**Supplementary Table 12c: Complete list of regression coefficients of models trained by postprandial metabolomics changes.**

**Supplementary Table 12d: Complete list of regression coefficients of models trained by microbial species.**

**Supplementary Table 12e: Complete list of regression coefficients of models trained by microbial KOs.**

**Supplementary Table 13: Associations of different medications with the gut microbiome, serum metabolome and transcriptome of different human tissues, evaluated by PERMANOVA analysis.**

## References

1. Alberti, K.G. & Zimmet, P.Z. Definition, diagnosis and classification of diabetes mellitus and its complications. Part 1: diagnosis and classification of diabetes mellitus provisional report of a WHO consultation. Diabet Med 15, 539–553 (1998).

2. Collaboration, N.C.D.R.F. Worldwide trends in diabetes since 1980: a pooled analysis of 751 population-based studies with 4.4 million participants. Lancet 387, 1513–1530 (2016).

3. Zimmet, P.Z. Diabetes and its drivers: the largest epidemic in human history? Clin Diabetes Endocrinol 3, 1 (2017).

4. Tabak, A.G., Herder, C., Rathmann, W., Brunner, E.J. & Kivimaki, M. Prediabetes: a high-risk state for diabetes development. Lancet 379, 2279–2290 (2012).

5. American Diabetes, A. Diagnosis and classification of diabetes mellitus. Diabetes Care 34 **Suppl 1**, S62–69 (2011).

6. Corpeleijn, E., Saris, W.H. & Blaak, E.E. Metabolic flexibility in the development of insulin resistance and type 2 diabetes: effects of lifestyle. Obes Rev 10, 178–193 (2009).

7. Goodpaster, B.H. & Sparks, L.M. Metabolic Flexibility in Health and Disease. Cell metabolism 25, 1027–1036 (2017).

8. DeFronzo, R.A. & Tripathy, D. Skeletal muscle insulin resistance is the primary defect in type 2 diabetes. Diabetes Care 32 **Suppl 2**, S157–163 (2009).

9. Cnop, M., et al. Mechanisms of pancreatic beta-cell death in type 1 and type 2 diabetes: many differences, few similarities. Diabetes 54 **Suppl 2**, S97–107 (2005).

10. Laiteerapong, N., et al. The Legacy Effect in Type 2 Diabetes: Impact of Early Glycemic Control on Future Complications (The Diabetes & Aging Study). Diabetes Care 42, 416–426 (2019).

11. Greenbaum, C.J., et al. Mixed-Meal Tolerance Test Versus Glucagon Stimulation Test for the Assessment of beta-Cell Function in Therapeutic Trials in Type 1 Diabetes. Diabetes Care 31, 1966–1971 (2008).

12. Ahmed, M., Gannon, M.C. & Nuttall, F.Q. Postprandial plasma glucose, insulin, glucagon and triglyceride responses to a standard diet in normal subjects. Diabetologia 12, 61–67 (1976).

13. Besser, R.E., Shields, B.M., Casas, R., Hattersley, A.T. & Ludvigsson, J. Lessons from the mixed-meal tolerance test: use of 90-minute and fasting C-peptide in pediatric diabetes. Diabetes Care 36, 195–201 (2013).

14. Besser, R.E.J., et al. The impact of insulin administration during the mixed meal tolerance test. Diabetic Med 29, 1279–1284 (2012).

15. Wopereis, S., et al. Multi-parameter comparison of a standardized mixed meal tolerance test in healthy and type 2 diabetic subjects: the PhenFlex challenge. Genes Nutr 12, 21 (2017).

16. Pellis, L., et al. Plasma metabolomics and proteomics profiling after a postprandial challenge reveal subtle diet effects on human metabolic status. Metabolomics 8, 347–359 (2012).

17. Shankar, S.S., et al. Standardized Mixed-Meal Tolerance and Arginine Stimulation Tests Provide Reproducible and Complementary Measures of beta-Cell Function: Results From the Foundation for the National Institutes of Health Biomarkers Consortium Investigative Series. Diabetes Care 39, 1602–1613 (2016).

18. Zeevi, D., et al. Personalized Nutrition by Prediction of Glycemic Responses. Cell 163, 1079–1094 (2015).

19. Berry, S.E., et al. Human postprandial responses to food and potential for precision nutrition. Nat Med 26, 964–973 (2020).

20. Zhou, W.Y., et al. Longitudinal multi-omics of host-microbe dynamics in prediabetes. Nature 569, 663-+ (2019).

21. Pedersen, H.K., et al. Human gut microbes impact host serum metabolome and insulin sensitivity. Nature 535, 376–381 (2016).

22. Backman, M., et al. Multi-omics insights into functional alterations of the liver in insulin-deficient diabetes mellitus. Mol Metab 26, 30–44 (2019).

23. Menni, C., et al. Serum metabolites reflecting gut microbiome alpha diversity predict type 2 diabetes. Gut microbes 11, 1632–1642 (2020).

24. Zhao, X.J., et al. Metabonomic fingerprints of fasting plasma and spot urine reveal human pre-diabetic metabolic traits. Metabolomics 6, 362–374 (2010).

25. Holecek, M. Branched-chain amino acids in health and disease: metabolism, alterations in blood plasma, and as supplements. Nutr Metab (Lond) 15, 33 (2018).

26. White, P.J. & Newgard, C.B. Branched-chain amino acids in disease. Science 363, 582–583 (2019).

27. Jenkinson, C.P., et al. Transcriptomics in type 2 diabetes: Bridging the gap between genotype and phenotype. Genom Data 8, 25–36 (2016).

28. Saxena, A., et al. Transcriptome profiling reveals association of peripheral adipose tissue pathology with type-2 diabetes in Asian Indians. Adipocyte 8, 125–136 (2019).

29. Fadista, J., et al. Global genomic and transcriptomic analysis of human pancreatic islets reveals novel genes influencing glucose metabolism. Proc Natl Acad Sci U S A 111, 13924–13929 (2014).

30. Mathur, S.K., et al. Transcriptomic analysis of visceral adipose from healthy and diabetic obese subjects. Indian J Endocrinol Metab 17, 446–450 (2013).

31. Thingholm, L.B., et al. Obese Individuals with and without Type 2 Diabetes Show Different Gut Microbial Functional Capacity and Composition. Cell Host & Microbe 26, 252-+ (2019).

32. Karlsson, F.H., et al. Gut metagenome in European women with normal, impaired and diabetic glucose control. Nature 498, 99–103 (2013).

33. Zhong, H., et al. Distinct gut metagenomics and metaproteomics signatures in prediabetics and treatment-naive type 2 diabetics. EBioMedicine 47, 373–383 (2019).

34. Gurung, M., et al. Role of gut microbiota in type 2 diabetes pathophysiology. EBioMedicine 51, 102590 (2020).

35. Wu, H., et al. The Gut Microbiota in Prediabetes and Diabetes: A Population-Based Cross-Sectional Study. Cell metabolism (2020).

36. Koh, A., et al. Microbially Produced Imidazole Propionate Impairs Insulin Signaling through mTORC1. Cell 175, 947–961 e917 (2018).

37. Van Olden, C.C., et al. A Systems Biology approach to understand gut microbiota and host metabolism in morbid obesity: design of the BARIA Longitudinal Cohort Study. Journal of internal medicine (2020).

38. American Diabetes, A. 2. Classification and Diagnosis of Diabetes: Standards of Medical Care in Diabetes-2020. Diabetes Care 43, S14–S31 (2020).

39. Yu, D., et al. Plasma metabolomic profiles in association with type 2 diabetes risk and prevalence in Chinese adults. Metabolomics 12(2016).

40. Lee, S., et al. Integrated Network Analysis Reveals an Association between Plasma Mannose Levels and Insulin Resistance. Cell metabolism 24, 172–184 (2016).

41. Ferrannini, E., et al. Mannose is an insulin-regulated metabolite reflecting whole-body insulin sensitivity in man. Metabolism 102, 153974 (2020).

42. Lu, Y., et al. Serum Amino Acids in Association with Prevalent and Incident Type 2 Diabetes in A Chinese Population. Metabolites 9(2019).

43. Mooradian, A.D. Dyslipidemia in type 2 diabetes mellitus. Nat Clin Pract Endocrinol Metab 5, 150–159 (2009).

44. Doumatey, A.P., et al. Gut Microbiome Profiles Are Associated With Type 2 Diabetes in Urban Africans. Front Cell Infect Microbiol 10, 63 (2020).

45. Newgard, C.B., et al. A branched-chain amino acid-related metabolic signature that differentiates obese and lean humans and contributes to insulin resistance. Cell metabolism 9, 311–326 (2009).

46. White, P.J., et al. The BCKDH Kinase and Phosphatase Integrate BCAA and Lipid Metabolism via Regulation of ATP-Citrate Lyase. Cell metabolism 27, 1281–1293 e1287 (2018).

47. Newgard, C.B. Interplay between lipids and branched-chain amino acids in development of insulin resistance. Cell metabolism 15, 606–614 (2012).

48. Hoyles, L., et al. Molecular phenomics and metagenomics of hepatic steatosis in non-diabetic obese women. Nat Med 24, 1070–1080 (2018).

49. Nemet, I., et al. A Cardiovascular Disease-Linked Gut Microbial Metabolite Acts via Adrenergic Receptors. Cell 180, 862–877 e822 (2020).

50. Witjes, J.J., Smits, L.P., Pekmez, C.T., Prodan, A. & Meijnikman, A.S. Donor Fecal Microbiota Transplantation Alters Gut Microbiota and Metabolites in Obese Individuals With Steatohepatitis. Hepatology communications (2020).

51. Jagadeeshaprasad, M.G., et al. Targeted quantification of N-1-(carboxymethyl) valine and N-1- (carboxyethyl) valine peptides of beta-hemoglobin for better diagnostics in diabetes. Clin Proteomics 13, 7 (2016).

52. Daniele, G., et al. Plasma N-Acetylaspartate Is Related to Age, Obesity, and Glucose Metabolism: Effects of Antidiabetic Treatment and Bariatric Surgery. Front Endocrinol (Lausanne) 11, 216 (2020).

53. Surendran, S. & Bhatnagar, M. Upregulation of N-acetylaspartic acid induces oxidative stress to contribute in disease pathophysiology. Int J Neurosci 121, 305–309 (2011).

54. Kashyap, S.R., et al. Insulin resistance is associated with impaired nitric oxide synthase activity in skeletal muscle of type 2 diabetic subjects. J Clin Endocrinol Metab 90, 1100–1105 (2005).

55. Tessari, P., et al. Nitric oxide synthesis is reduced in subjects with type 2 diabetes and nephropathy. Diabetes 59, 2152–2159 (2010).

56. Rizzo, N.O., et al. Reduced NO-cGMP signaling contributes to vascular inflammation and insulin resistance induced by high-fat feeding. Arterioscler Thromb Vasc Biol 30, 758–765 (2010).

57. Al-Sulaiti, H., et al. Metabolic signature of obesity-associated insulin resistance and type 2 diabetes. J Transl Med 17, 348 (2019).

58. Shah, P., et al. Elevated free fatty acids impair glucose metabolism in women: decreased stimulation of muscle glucose uptake and suppression of splanchnic glucose production during combined hyperinsulinemia and hyperglycemia. Diabetes 52, 38–42 (2003).

59. Roden, M., et al. Mechanism of free fatty acid-induced insulin resistance in humans. J Clin Invest 97, 2859–2865 (1996).

60. Ruiz de Azua, I., Gautam, D., Jain, S., Guettier, J.M. & Wess, J. Critical metabolic roles of beta-cell M3 muscarinic acetylcholine receptors. Life Sci 91, 986–991 (2012).

61. Gautam, D., et al. A critical role for beta cell M3 muscarinic acetylcholine receptors in regulating insulin release and blood glucose homeostasis in vivo. Cell metabolism 3, 449–461 (2006).

62. Zhang, Y., et al. Regulation of Adipose Tissue Inflammation and Insulin Resistance by MAPK Phosphatase 5. J Biol Chem 290, 14875–14883 (2015).

63. Cawthorn, W.P. & Sethi, J.K. TNF-alpha and adipocyte biology. FEBS Lett 582, 117–131 (2008).

64. Hotamisligil, G.S., Shargill, N.S. & Spiegelman, B.M. Adipose expression of tumor necrosis factor-alpha: direct role in obesity-linked insulin resistance. Science 259, 87–91 (1993).

65. Palmer, A.K., et al. Cellular Senescence in Type 2 Diabetes: A Therapeutic Opportunity. Diabetes 64, 2289–2298 (2015).

66. Santulli, G., et al. Calcium release channel RyR2 regulates insulin release and glucose homeostasis. J Clin Invest 125, 1968–1978 (2015).

67. Morton, J.T., et al. Learning representations of microbe-metabolite interactions. Nat Methods 16, 1306–1314 (2019).

68. Albareda, M., Rodriguez-Espinosa, J., Murugo, M., de Leiva, A. & Corcoy, R. Assessment of insulin sensitivity and beta-cell function from measurements in the fasting state and during an oral glucose tolerance test. Diabetologia 43, 1507–1511 (2000).

69. Matthews, D.R., et al. Homeostasis model assessment: insulin resistance and beta-cell function from fasting plasma glucose and insulin concentrations in man. Diabetologia 28, 412–419 (1985).

70. Wallace, T.M., Levy, J.C. & Matthews, D.R. Use and abuse of HOMA modeling. Diabetes Care 27, 1487–1495 (2004).

71. Brouns, F., et al. Glycaemic index methodology. Nutr Res Rev 18, 145–171 (2005).

72. Brodovicz, K.G., et al. Postprandial metabolic responses to mixed versus liquid meal tests in healthy men and men with type 2 diabetes. Diabetes Res Clin Pract 94, 449–455 (2011).

73. Tyanova, S., et al. The Perseus computational platform for comprehensive analysis of (prote)omics data. Nat Methods 13, 731–740 (2016).

74. Ewels, P.A., et al. The nf-core framework for community-curated bioinformatics pipelines. Nature biotechnology 38, 276–278 (2020).

75. Ewels, P., Magnusson, M., Lundin, S. & Kaller, M. MultiQC: summarize analysis results for multiple tools and samples in a single report. Bioinformatics 32, 3047–3048 (2016).

76. Dobin, A., et al. STAR: ultrafast universal RNA-seq aligner. Bioinformatics 29, 15–21 (2013).

77. Liao, Y., Smyth, G.K. & Shi, W. featureCounts: an efficient general purpose program for assigning sequence reads to genomic features. Bioinformatics 30, 923–930 (2014).

78. Pertea, M., et al. StringTie enables improved reconstruction of a transcriptome from RNA-seq reads. Nature biotechnology 33, 290–295 (2015).

79. Di Tommaso, P., et al. Nextflow enables reproducible computational workflows. Nature biotechnology 35, 316–319 (2017).

80. Love, M.I., Huber, W. & Anders, S. Moderated estimation of fold change and dispersion for RNA-seq data with DESeq2. Genome Biol 15, 550 (2014).

81. Varemo, L., Nielsen, J. & Nookaew, I. Enriching the gene set analysis of genome-wide data by incorporating directionality of gene expression and combining statistical hypotheses and methods. Nucleic Acids Res 41, 4378–4391 (2013).

82. Costea, P.I., et al. Towards standards for human fecal sample processing in metagenomic studies. Nature biotechnology 35, 1069–1076 (2017).

83. Karlsson, F.H., Nookaew, I. & Nielsen, J. Metagenomic data utilization and analysis (MEDUSA) and construction of a global gut microbial gene catalogue. PLoS Comput Biol 10, e1003706 (2014).

84. Langmead, B. & Salzberg, S.L. Fast gapped-read alignment with Bowtie 2. Nat Methods 9, 357–359 (2012).

85. Dixon & Philip. VEGAN, a package of R functions for community ecology. Journal of Vegetation Science 14, 927–930 (2003).

86. Thioulouse, J., Dray, S. & Dufour, A.-B. *Multivariate Analysis of Ecological Data with ade4*, (2018).

87. Friedman, J., Hastie, T. & Tibshirani, R. Regularization Paths for Generalized Linear Models via Coordinate Descent. J Stat Softw 33, 1–22 (2010).

